# Foxp1 acts upstream of Vegfa, suppresses cortical angiogenesis, and promotes hypoxia in radial glia

**DOI:** 10.1101/2022.10.17.512545

**Authors:** Caroline A. Pearson, Jessie E. Buth, Michael R.M. Harrison, M. Elizabeth Ross, Bennett G. Novitch

## Abstract

Radial glia progenitors within the cerebral cortex undergo a characteristic switch between symmetric self-renewing cell divisions early in development and asymmetric neurogenic divisions at later times, yet the mechanisms controlling this transition remain unclear. Previous work has shown that the autism-linked transcription factor Foxp1 is endogenously expressed by early but not late radial glia, and both loss and gain of Foxp1 can alter their neural progenitor activities and fate choices. Here, we show that premature loss of Foxp1 leads to an increase in transcriptional programs regulating angiogenesis, glycolysis, and cellular responses to hypoxia. These changes coincide with an elevation in Vegfa expression in radial glia and precocious vascular network development. Thus, the endogenous decline in Foxp1 expression appears to orchestrate changes in the tissue environment adjacent to radial glia that influence their metabolic state which in turn can alter their self-renewal and neurogenic capacities.

## INTRODUCTION

The temporal specification of neuronal subtypes in the mammalian cortex arises through progressive changes in the transcriptional states of cortical progenitors termed radial glia (RG) (Desai and McConnell, 2000; Shen et al., 2006; Telley et al., 2019). As neurogenesis proceeds, RG potential becomes more restricted and increasingly influenced by extrinsic factors and substrates (Desai and McConnell, 2000; Heavner et al., 2020; Magrinelli et al., 2022; Shen *et al*., 2006; Telley *et al*., 2019). RG line the ventricular zone (VZ), proliferate and self-renew to maintain a progenitor population. Early RG (~embryonic day E12-13 in mouse) have the potential to generate all classes of excitatory glutamatergic neurons and glia (Beattie and Hippenmeyer, 2017; Rakic, 2003). Late RG (~E13.5-15.5 in mouse) generate basal progenitors including intermediate progenitors and basal radial glia that, in turn, amplify the generation of later-born excitatory neurons which make up the upper layers of the cortex (Beattie and Hippenmeyer, 2017; Molnár et al., 2019; Paridaen and Huttner, 2014; Taverna et al., 2014). The precise timing of the switch from early multipotent RG to late more restricted RG is integral to the layered organization of cortex and orderly assembly of cortical circuits.

We previously demonstrated that the autism-linked transcription factor Foxp1 is highly expressed by early RG, promotes their symmetric cell divisions and self-renewal, and sustains their broad potential to generate both early-born deep layer neurons and later-born superficial layer neurons (Pearson et al., 2020). The endogenous downregulation of Foxp1 in late RG at mid-neurogenic stages coincides with and is required for the transition to asymmetric neurogenic divisions. Conditional removal of Foxp1 function from early RG resulted in a premature transition towards later characteristics including increased basal progenitor generation and neuronal differentiation, resulting in a reduction of early-born deep layer neurons and a concomitant increase in upper layer neurons (Pearson *et al*., 2020). However, the mechanisms through which Foxp1 promotes early RG character are unclear.

Recent single cell RNA-Seq studies have shown that RG become increasingly responsive to extrinsic signals in the embryonic environment as development proceeds, particularly substrates made available through the developing vascular network (Dong et al., 2022; Telley *et al*., 2019). Two of the major substrates provided include oxygen and glucose, which are vital for energy production via lactate production and the tricarboxylic acid cycle. The switch of RG from symmetric self-renewing cell divisions to asymmetric neurogenic divisions has been associated with the relief from hypoxia as the vascular network forms and increases in RG glycolytic activity (Javaherian and Kriegstein, 2009; Komabayashi-Suzuki et al., 2019; Panchision, 2009; Wagenführ et al., 2015). Nevertheless, the developmental factors that coordinate angiogenesis with neurogenesis remain poorly defined.

In this study, we show that early loss of Foxp1 in RG leads to immediate transcriptional changes in genes associated with angiogenesis, Hypoxia Inducible factor 1 alpha (HIF1α) signaling, and glycolysis. In situ hybridization (ISH) and immunohistochemical (IHC) analyses demonstrated many misregulated glycolysis genes are specifically expressed by RG. Furthermore, one of the most upregulated genes, Vascular endothelial growth factor A (Vegfa) is upregulated in RG as Foxp1 levels decline. These changes coincide with the premature development of vasculature. Collectively, these findings reveal that Foxp1 acts upstream of angiogenic and glycolytic programs, coupling the timing of blood vessel development with the transition from early to late RG character.

## RESULTS

### The endogenous downregulation of Foxp1 coincides with the establishment of the cortical vasculature

To begin to understand the role of Foxp1 in directing the transition from self-renewal to neurogenic divisions, we asked whether the endogenous downregulation of Foxp1 coincided with changes in cortical vasculature and the switch to intermediate progenitor generation. IHC analysis of Foxp1 protein levels in RG and the presence of laminin^+^ blood vessels confirmed that at E12.5, when Foxp1 levels are highest, there are few laminin^+^ blood vessels in the lateral cortex (Figures 1A, 1B, 1I, and 1J). Downregulation of Foxp1 at E13.5-E14.5 coincides with an increase in the number of vessels (Figures 1C, 1D, 1I, and 1J). Analysis of Tbr2^+^ intermediate progenitors demonstrated a concomitant increase in these cells from E13.5 onwards (Figures 1E–1H and 1K). Therefore, the downregulation of Foxp1 is coincident with the increased vascularization of the cortex and the generation of intermediate progenitors.

**Figure 1.**
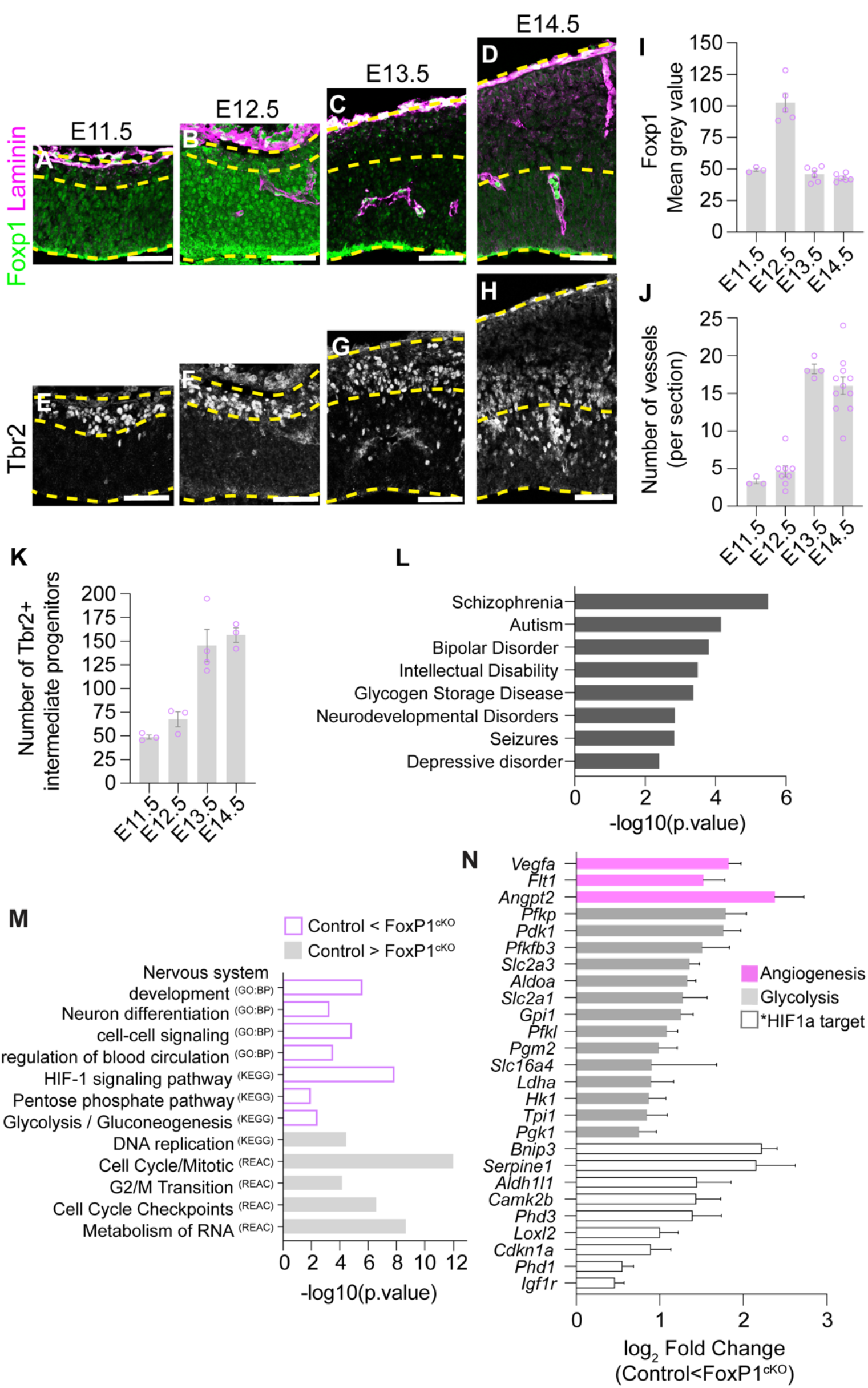
Foxp1 downregulation coincides with the expansion of the cortical vasculature and transcriptomic changes associated with angiogenesis, glycolysis and HIF signaling. (A-D) Laminin and Foxp1 protein expression in the lateral cortex from E11.5 to E14.5. (E-H) IHC for Tbr2 in the lateral cortex from E11.5-E14.5. (I) Quantification of Foxp1 mean grey value in RG between E11.5 and E14.5. (J) Quantification of laminin^+^ PVP vessels per section of the lateral cortex between E11.5-E14.5. (K) Quantification of Tbr2^+^ intermediate progenitors in the lateral cortex from E11.5-E14.5. (L) Human disorders associated with genes significantly misregulated in Foxp1^cKO^ mutants at E12.5. (M) Gene ontology analysis of terms associated with significantly misregulated genes in Foxp1^cKO^ lateral cortex at E12.5. GO-BP, biological process; KEGG, pathways, REAC, reactome pathway. (N) Significantly misregulated genes associated with angiogenesis (magenta), glycolysis (grey) and HIF1α signaling (white) in Foxp1^cKOs^ at E12.5. *denotes genes that are also HIF1α target genes. Measurements from 3-5 sections per embryo, 3 embryos per time point. Scale bars 50um.

### Foxp1 loss leads to the upregulation of genes associated with neuronal differentiation, angiogenesis, HIF-1 signaling, and glycolysis in early radial glia

To determine the transcriptional changes accompanying early Foxp1 loss, we performed bulk RNA-Seq analysis comparing control *(Emx1^Cre^* negative littermates) and *Foxp1^fl/fl^; Emx1^Cre/+^* (termed *Foxp1^cKO^*) mutant lateral cortices collected at E12.5 (a time point at which the majority of cells are RG). Of the 514 significantly misregulated genes, 307 were upregulated (log2 fold change >0.5) and 80 were downregulated (log_2_ fold change <-0.5), consistent with the reported role of Foxp1 as a transcriptional repressor in other tissues (Zhang et al., 2010) (Figure S1A, Tables S2-S3). The main human disease categories associated with genes upregulated in the absence of Foxp1 included schizophrenia, Autism, intellectual disorders, neurodevelopmental disorders, glycogen storage disorders, and seizures (Figure 1L). Gene ontology analyses showed that terms associated with biological processes such as nervous system development, neuron differentiation and cellcell signaling were overrepresented in the absence of Foxp1, whilst terms associated with DNA replication, cell cycle/mitosis and RNA metabolism were underrepresented (Figure 1M). Amongst the most overrepresented processes and pathways in the absence of Foxp1 were responses to cellcell signaling, regulation of blood circulation, glycolysis/gluconeogenesis, and HIF-1 signaling pathway (Figure 1M). Consistent with these findings, we found that many key genes involved in glycolysis, HIF-1 signaling, and angiogenesis were upregulated in the absence of *Foxp1* including *Vegfa, Ldha, Slc2a1* and *Phd2* (Figure 1N). Thus, early loss of Foxp1 in RG leads to transcriptional increases in genes associated with differentiation, angiogenesis, and increased dependence on extrinsic factors such as glucose and oxygen.

### Genes associated with angiogenesis, glycolysis and HIF1α signaling are specifically expressed by early RG

Due to the heterogeneity of cell types in our RNA-Seq samples, we sought to determine which of the observed transcriptional changes were intrinsic to RG. To demonstrate the regional expression of misregulated genes associated with cell cycle, neurogenesis, angiogenesis, glycolysis, and HIF1α signaling, we performed ISH in wildtype tissue at E12.5. This analysis confirmed the predicted expression of neuronal genes, *Camk2b, Dlg4, Neo1* and *Sez6* transcripts in the nascent cortical plate (Figures S2A-S2D) and cell cycle gene *Ccne1* in the VZ (Figure S2E). Several misregulated genes involved in glycolysis, HIF signaling, and angiogenesis were specifically expressed in RG at E12.5, suggesting cell autonomous effects of Foxp1 loss in this population. Glycolysis genes specifically expressed in the VZ included *Aldoa, Ldha, Slc16a4* (encodes the lactate transporter Monocarboxylate transporter 4), *Pfkl, Pfkfb3*, and *Pdk1* (Figures 2A, 2B, and S2F-S2J). IHC analysis of Ldha with Nestin confirmed its association with RG (Figures 2C and 2D). Expression of HIF1α target genes *Phd2* and *Loxl2* were also detected in the VZ (Figures 2E and 2F). Glucose transporter 1 (Glut1 encoded by *Slc2a1)* is also expressed in the VZ as well as endothelial cells (Figure 2G). IHC analysis showed that Glut1 protein is expressed in Isolectin B4^+^ (IB4) periventricular plexus (PVP) endothelial cells (indicated by white asterisks in Figure 2H) and in Nestin^+^ RG with pronounced accumulation at their apical endfeet (Figures 2H and 2I, indicated by white arrowheads in 2H).

**Figure 2.**
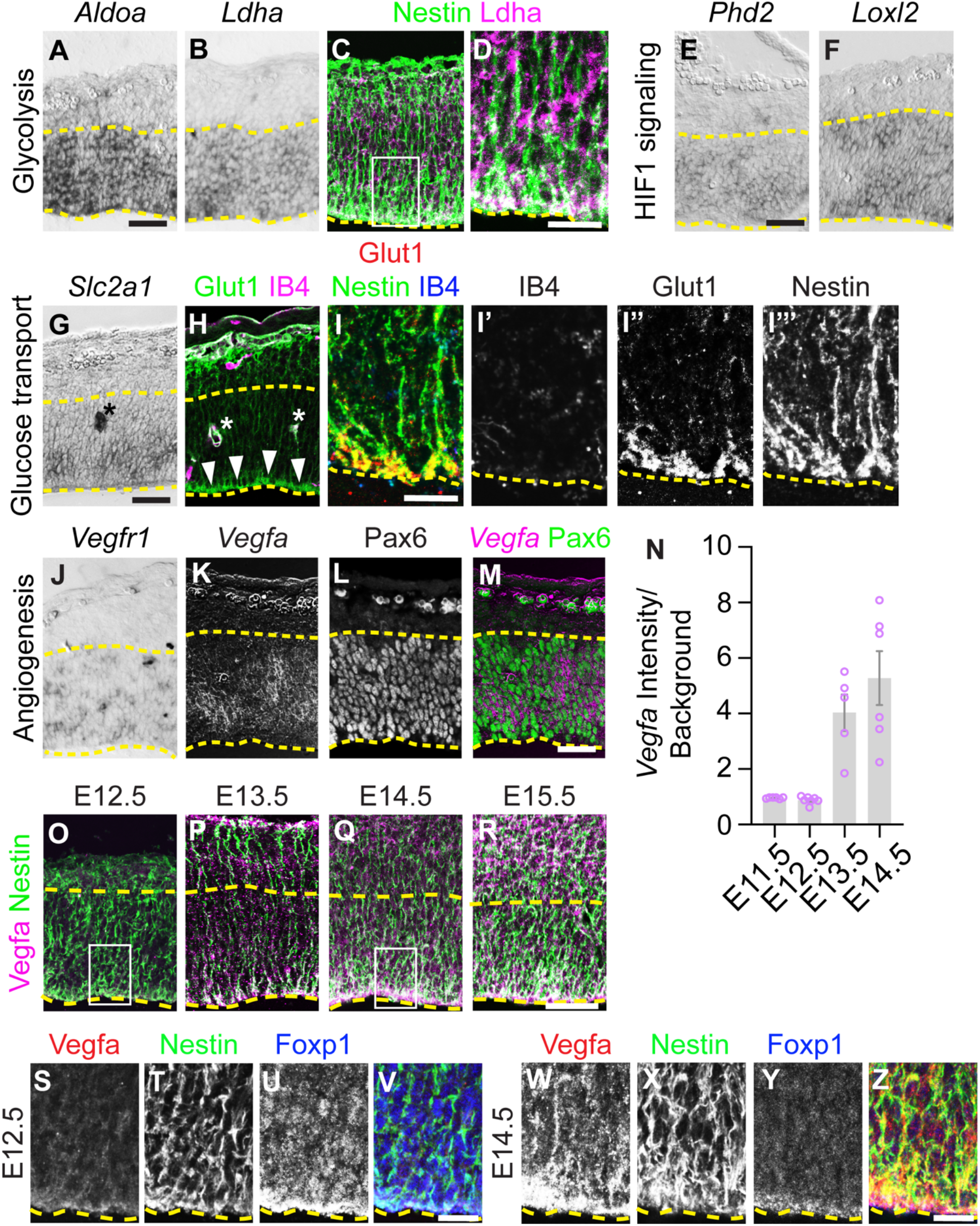
Gene expression associated with early radial glia in the wildtype cortex. (A-B) mRNA expression of glycolysis genes *Aldoa, Ldha* in the lateral cortex at E12.5. (C-D) Ldha and Nestin protein expression in the lateral cortex at E12.5. Boxed area in C denotes magnified area in D. (E-F) mRNA expression of HIF1α target genes *Phd2* and Loxl2 in the VZ at E12.5. (G) *Slc2a1* mRNA expression in the VZ at E12.5. (H) IHC analysis of Glut1 and Isolectin B4 (IB4) at E12.5. Arrowheads highlight apical accumulation. (I) IHC for Glut1, IB4 and Nestin at E12.5. I’-I’’’’ show single channel images. (J) Vegfr1 mRNA expression in the lateral cortex at E12.5. (K-M) *Vegfa* mRNA expression in Pax6^+^ RG in the VZ at E12.5. (N) Quantification of *Vegfa* intensity (over background) in the lateral cortex from E11.5-E14.5. (O-R) IHC for Vegfa and Nestin between E12.5 and E15.5 in the lateral cortex. (S-Z) IHC for Vegfa, Nestin and Foxp1 in the VZ at E12.5 and E14.5. S-V is high magnification of boxed area in O. W-Z is high magnification of boxed area in Q. Yellow dashed lines demarcate the VZ. Scale bars 50μm (A-M, O-R), 20μm (S-Z). Measurements from 2-3 sections per embryo, 5-6 embryos per time point.

Other genes identified in our RNA Seq analyses were not specifically expressed by RG, suggesting non-cell autonomous effects of Foxp1 loss on other cell populations. For instance, Glucose transporter 3 *(Slc2a3* encodes Glut3) expression was not detected in the VZ but was instead detected in pial perineural plexus (PNVP) endothelial cells (Figure S2I). Endothelial cell specific expression of *Cdh13* and *Igfbp3* was also detected (Figures S2K and S2L). Vegf receptor 1 *(Vegfr1)* expression was detected in PNVP and periventricular plexus (PVP) endothelial cells at high levels, and at lower levels in a salt and pepper fashion in the VZ (Figure 2J). A similar expression pattern was seen by RNAScope fluorescence ISH at E13.5 (Figure S2M).

*Vegfa* was identified as one of the most upregulated genes in our RNA-Seq analysis (Figure S1A). Previous studies have demonstrated that Vegfa protein is a critical regulator of cortical vascularization (Haigh et al., 2003; Mackenzie and Ruhrberg, 2012; Raab et al., 2004; Tata et al., 2015). At E12.5, low levels of *Vegfa* are expressed in Pax6^+^ RG (Figure 2K–2M). ISH analysis demonstrated an upregulation of *Vegfa* expression between E12.5 to E13.5, followed by a further increase at E14.5 (Figures 2N and S3A-S3D). This trend was confirmed by RNAScope analysis of *Vegfa* which showed that it is expressed in the VZ at E13.5 and E15.5 (Figures S3E and S3F). Comparison of expression levels in the VZ confirmed a further increase in *Vegfa* expression at E15.5 (Figure S3I). At this later timepoint, *Vegfa* expression was also detected in the intermediate zone and cortical plate, albeit at lower levels than that found in the VZ (Figure S3F). IHC analysis of Vegfa protein between E12.5 and E15.5 demonstrated its upregulation in Nestin^+^ RG from E13.5 onwards, with accumulation at the apical surface of the VZ (Figure 2O-R, Figures S3G, S3 H, S3J-S3O). Analysis of Vegfa with the RG marker Nestin and Foxp1 demonstrated that at E12.5 Nestin^+^ RG express high levels of Foxp1 and low levels of Vegfa (Figure 3S–3V). By contrast, E14.5, Nestin^+^ RG express low levels of Foxp1 and increased levels of Vegfa (Figures 2W–2Z). Thus, Vegfa is expressed by RG and its upregulation coincides with the endogenous downregulation of Foxp1.

**Figure 3.**
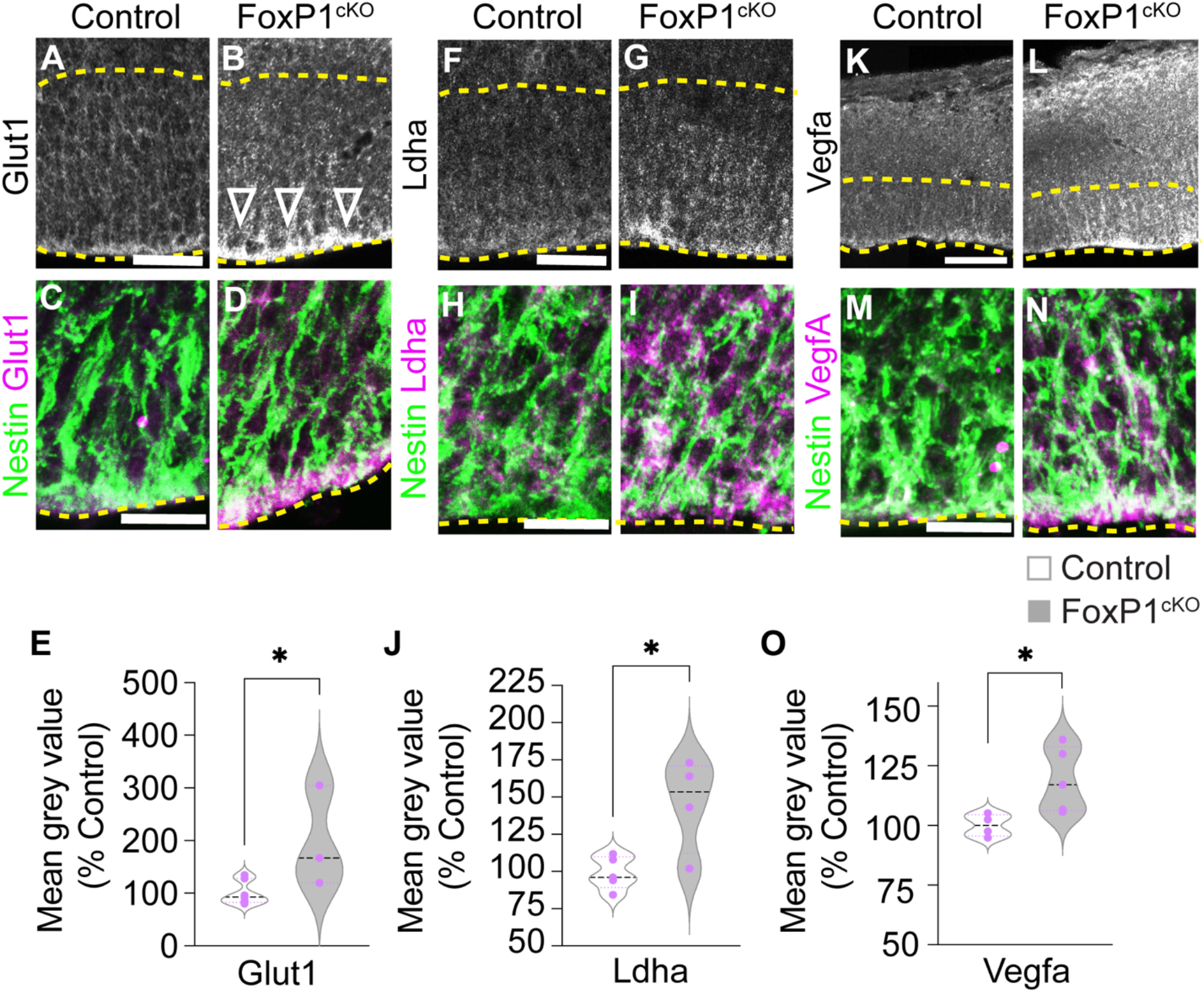
Loss of Foxp1 results in upregulation of HIF1α targets including Vegfa protein. (A-B) IHC for Glut1 in control and Foxp1^cKO^ mutants in the VZ of the lateral cortex at E12.5. (CD) Overlap between Glut1 and Nestin expression in control and Foxp1^cKO^ mutants. (E) Quantification of Glut1 mean grey value (percent control) in Control and Foxp1^cKO^ mutant lateral cortex at E12.5. N=2 litters, 5 controls, 4 mutants, p*=0.0262. (F-G) IHC for Ldha in control and Foxp1^cOn^ mutants in the lateral cortex at E12.5. (H-I) Overlap between Ldha and Nestin expression in control and Foxp1^cKO^ mutants. (J) Quantification of Ldha mean grey value (percent control) in Control and Foxp1^cKO^ mutant lateral cortex at E12.5. N=2 litters, 5 controls, 4 mutants, p*=.0169. (K-L) IHC for Vegfa in control and Foxp1^cKO^ mutants in the VZ of the lateral cortex at E12.5. (MN) Overlap between Vegfa and Nestin expression in control and Foxp1^cKO^ mutants. (O) Quantification of Vegfa mean grey value (percent control) in Control and Foxp1^cOn^ lateral cortex at E12.5. N= 2 litters, 4 controls, 5 mutants, *p=0.0328. Scale bars 50μm (A, B, F, G), 20μm (C, D, H, I, M, N), 100μm (K-L). Dashed yellow lines demarcate VZ.

### Early loss of Foxp1 promotes glycolysis and early relief from hypoxia in RG

Our transcriptional analysis suggests that in the absence of Foxp1 there is an increase in RG glycolysis and activation of HIF1α target genes. We sought to confirm that genes associated with these processes are upregulated in RG in the absence of Foxp1. We selected Glut1 and Ldha as they are key components of the glycolysis pathway and known HIF1α targets. IHC analysis of Glut1 and Ldha in E12.5 control and Foxp1^cKO^ lateral cortices demonstrated that both proteins were significantly upregulated in the absence of Foxp1 and present in Nestin^+^ RG cells (Figures 3A–3J). We further confirmed upregulation of *Slc2a1* (encodes Glut1) and *Ldha* mRNA in the lateral cortex of Foxp1^cKO^ animals by qPCR (Figure S4D).

Upregulation of *Vegfa* mRNA in the E12.5 Foxp1cKO cortex was also confirmed by qPCR (Figure S4D) and increased abundance of Vegfa protein in Nestin^+^ RG cells by IHC (Figures 3K–3O). Next, we asked whether sustained expression of Foxp1 in the cortex is sufficient to repress Vegfa expression at mid-neurogenic stages. We used conditional Foxp1 transgenic mice to selectively activate Foxp1 expression in cortical RG using Emx::1Cre, termed Foxp1^cON^ (Pearson *et al*.,2020). Vegfa protein levels appeared slightly reduced in E14.5 Foxp1^cON^ cortices, but quantification did not surpass the threshold for statistical significance (p = 0.0877) (Figures S4A-S4C).

Together, these results suggest that Foxp1 expression in early RG is required to repress Vegfa, but its forced expression is not alone sufficient to repress Vegfa at later time points. The loss of Foxp1 also results in increased expression of HIF1α targets/glycolysis genes at the mRNA and protein level, indicating increased glycolysis and premature relief from hypoxia in the early cortex.

### Foxp1 repression of Vegfa signaling delays the development of the cortical vasculature

Given the upregulation of Vegfa in the absence of Foxp1, we examined how this manipulation impacted the development of the cortical vasculature. Using the endothelial cell marker IB4 in 50μm thick cryosections, we analyzed the ventral to dorsal expansion of the PVP network within the lateral cortex at E12.5 (Figures 4A–4D). At this stage, comparable numbers of vessels were present in the mid and ventral lateral cortex in control and Foxp1^cKO^ cortices (Figures 4A–4D and 4F). However, there appeared to be increased numbers of vessels in the dorsal lateral cortex in Foxp1^cKO^ embryos. To quantify the ventral-dorsal establishment of the PVP vasculature, we divided each section into 3 areas of equal size (Figure 4E) and counted the number of vessels in each region which confirmed a significant increase in the number of vessels in the dorsal-most lateral cortex in Foxp1^cKO^ embryos (Figure 4F). We further observed an increase in the number of tip cell filopodia in Foxp1^cKO^ samples at E12.5 compared to controls (Figures 4G–4I), which is consistent with previous observations that increased Vegfa signaling can induce the formation of endothelial tip cell filopodia (Gerhardt et al., 2003; Haigh *et al*., 2003).

**Figure 4.**
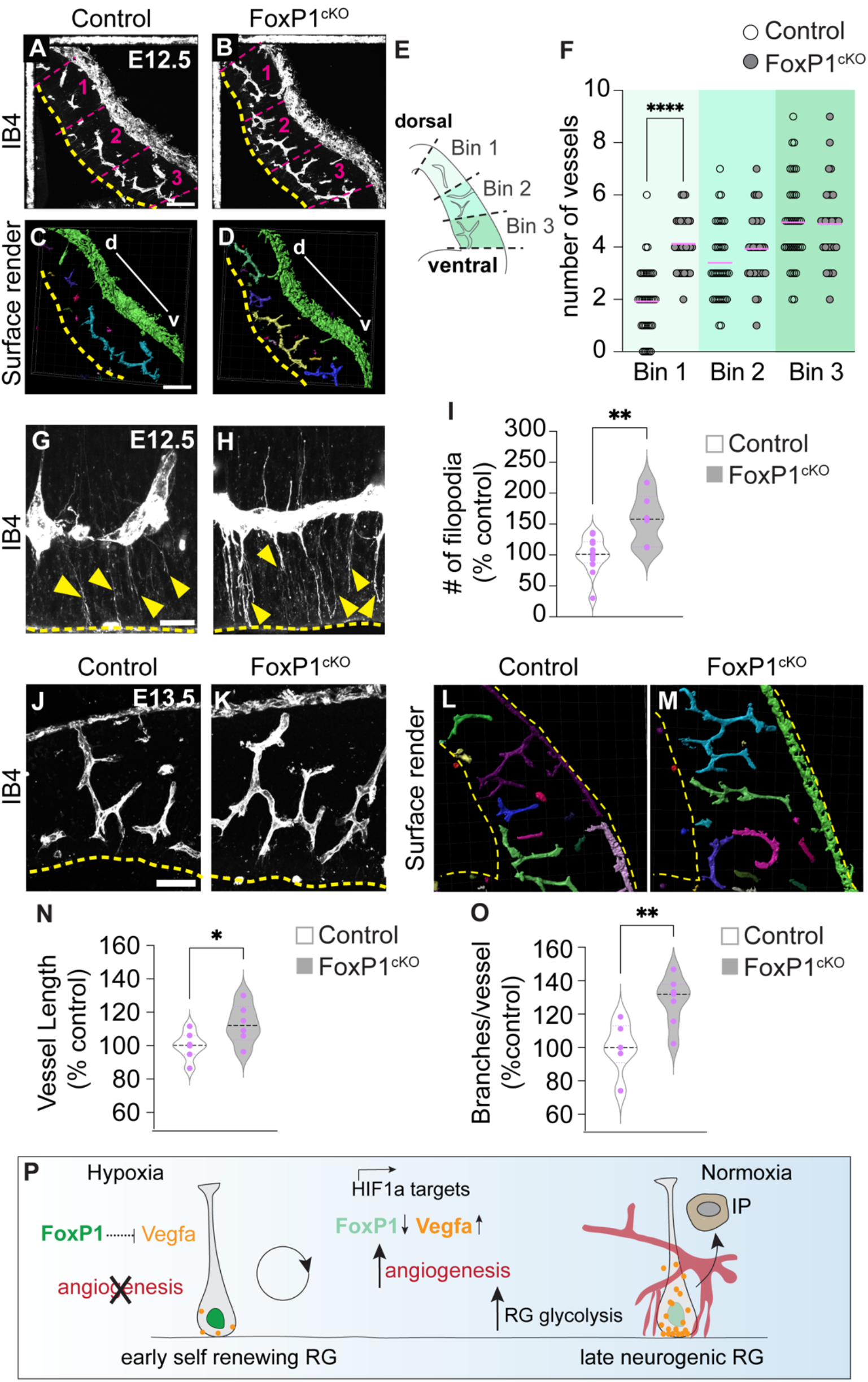
Loss of Foxp1 results in precocious development of the cortical vasculature. (A-B) Maximum intensity projection images of IB4 vessel staining in the lateral cortex of control and Foxp1^cKO^ embryos at E12.5. (C-D) Surface rendered images of A-B. Contiguous vessels are individually colored. (E) Schematic of lateral cortex divisions used for quantification in F. (F) Quantification of IB4^+^ vessels in each binned area in control and Foxp1^cKO^ lateral cortex at E12.5. N=3 litters, 7 mutants, 10 controls. Mann Whitney test ****p=<0.0001. (G-H) IB4^+^ filopodia at the apical surface (dotted line) at E12.5 in control and Foxp1^cKO^ cortex. Yellow arrowheads indicate filopodia. (I) Quantification of filopodia in control and Foxp1^cKO^ cortex. N = 3 litters, 7 mutants, 10 controls, **p=0.0033. (J-K) IB4^+^ blood vessels in control and Foxp1^cKO^ lateral cortex at E13.5. (L-M) Surface rendered images of lateral cortex in control and Foxp1^cKO^ at E13.5. Contiguous vessels are individually colored. (N-O) Quantification of vessel length and number of branches per vessel in control and Foxp1^cKO^ cortex. N= 3 litters, 8 mutants, 8 controls, *p=0.0386, **p=0.0063. (P) Schematic of Foxp1 mediated repression of cortical angiogenesis. Dashed yellow lines demarcate apical surface. d, dorsal; v, ventral. Scale bars 100μm (A-D), 20μm (G, H) 50μm (J, K).

We lastly analyzed the cortical vasculature a day later at E13.5. At this stage in control embryos, the lateral cortex has been perfused by vessels along the dorsoventral axis that have begun to form branches (Figures 4J and 4L). Using IB4 staining, we visualized blood vessels in control and Foxp1^cKO^ cortices and generated surface rendered images to identify contiguous vessels (Figures 4J–4M). Subsequent analyses of vessel length and morphology demonstrated that the vessels in Foxp1^cKO^ cortices are significantly longer and more branched compared to littermate controls (Figures 4N and 4O).

Collectively, these results demonstrate that Foxp1 expression in RG regulates the timing of cortical vascularization. Its absence results in an early establishment of cortical vasculature in the dorsal-most lateral cortex and precocious development of the vascular network in a non-stereotypical fashion.

## DISCUSSION

Neural stem cells in the developing cortex exhibit a tightly regulated progression in their modes of cell division and fate decisions which underpins the emergence of layered organization and subsequent circuit assembly. Our prior and current data collectively demonstrate that Foxp1 plays a key role in this process, gating the transition of RG through early-to-late stages of neurogenesis in part by modifying the stem cell niche microenvironment. Foxp1 acts upstream of Vegfa to suppress angiogenesis and sustain a hypoxic environment favorable for the early symmetric self-renewing cell divisions needed to expand the cortical progenitor pool (Figure 4P). As Foxp1 levels in RG decline over time, Vegfa production increases, facilitating the ingrowth of blood vessels and efficient delivery of oxygen and metabolites into the developing cortex. These events coincide with changes in the metabolic profile of the RG that favor asymmetric neurogenic cell divisions. Our findings complement and extend previous and new reports demonstrating that hypoxic conditions and the suppression of vascular ingrowth can repress the switch to neurogenesis (Dong *et al*., 2022; Lange et al., 2016).

Previous studies demonstrated that Vegfa expressed by RG forms a gradient that promotes angiogenesis (Haigh *et al*., 2003; Raab *et al*., 2004) and directs the migration of endothelial cells into the cortex (Tata and Ruhrberg, 2018). Vegfa induces endothelial filopodia formation in tip cells which direct sprouting angiogenesis and the extension of the vasculature. More recently a study elegantly revealed a reciprocal interaction between endothelial filopodia and mitotic RG in the ventral telencephalon. Mitotic RG induce endothelial filopodia growth by upregulating VegfA and, in turn, high endothelial filopodia density prolongs RG mitosis, favoring neuronal differentiation (Di Marco et al., 2020). Another study characterized the association of neural progenitors and vasculature in the developing cortex, revealing a similar physical interaction between dividing RG and endothelial filopodia. These studies highlight the interrelationship of the developing nervous and vascular systems as well as the importance of the nascent vasculature for cortical development (Komabayashi-Suzuki *et al*., 2019).

Other signals have been shown to regulate cortical angiogenesis, including Wnt signaling and Ldha (Daneman et al., 2009; Dong *et al*., 2022). Wnt genes were not identified as significantly misregulated in our RNA Seq analyses, suggesting Foxp1 acts independently of Wnt signaling to influence angiogenesis. However, the decreased vascularization we observe in the dorsal lateral cortex is of interest as different Wnt signals have been shown to regulate vascularization in the dorsal vs. ventral cortex (Daneman *et al*., 2009). The changes we see in the dorsal cortex could be due to early development and migration of the developing vascular network, or an earlier impact upon the Wnt signals that mediate angiogenesis in the dorsal cortex. Recently, Ldha has been shown to influence early stages of cortical angiogenesis, prior to the stages we have studied (Dong *et al*., 2022), and may continue to act in late RG. The changes seen in Ldha and other glycolysis genes in the absence of Foxp1 could reflect the increased availability of glucose in the RG microenvironment. Additionally, many glycolysis genes are HIF1α targets, and their upregulation indicates a transition from hypoxic to normoxic conditions. Further studies are required to determine whether aerobic or anaerobic glycolysis is dominant in Foxp1-deficient RG.

Lastly, an increasing body of evidence has implicated blood vessel pathologies in several neurodevelopmental disorders (Baruah and Vasudevan, 2019; Ouellette and Lacoste, 2021). For example, early defects in PVP endothelial cells have been linked to the origin of Autism (Azmitia et al., 2016). Given that *FOXP1* has been recognized as a high-confidence autism risk gene in humans, our findings raise the possibility that a significant impact of its mutation on human brain could be through its influence on the timing of angiogenesis and changes in the metabolic state of RG at the critical early stages of cortical development.

## STAR METHODS

### RESOURCE AVAILABILITY

#### Lead Contact

Further information and requests for resources and reagents should be directed to and will be fulfilled by the lead contact, Caroline Alayne Pearson.

#### Materials Availability

This study did not generate new unique reagents.

#### Data and Code Availability

- RNA-seq data have been deposited at NCBI Gene Expression Omnibus, GEO accession number GSE217364 and are publicly available as of the date of publication. Accession numbers are listed in the key resources table. Microscopy data reported in this paper will be shared by the lead contact upon request.
- No original code was created.
- Any additional information required to reanalyze the data reported in this paper is available from the lead contact upon request.

### EXPERIMENTAL MODEL AND SUBJECT DETAILS

#### Mouse lines

Foxp1^flox/flox^, Foxp1^tg/+^ and Emx1^*Cre*^ mice were maintained as previously described (Feng et al., 2010; Gorski et al., 2002; Iwasato et al., 2004; Wang et al., 2004) following UCLA Chancellor’s Animal Research Committee husbandry guideline. Male and female embryos between embryonic day 10.5-15.5 were used in this study. Cre negative litter mates were used as controls.

### METHOD DETAILS

#### Tissue preparation

Embryonic cortices were fixed in 4% paraformaldehyde in PBS overnight. Tissues were cryosectioned (10-50 μm sections) and processed for immunohistochemistry, in situ hybridization, or RNA Scope fluorescent in situ hybridization as previously described (Pearson *et al*., 2020; Rousso et al., 2012).

#### RNA sample collection

E12.5 lateral cortex samples were lysed in QIAzol reagents and RNA was extracted following manufacturer’s instructions (miRNeasy Micro Kit, Qiagen). Six samples were used for RNA sequencing consisting of four control females and two Foxp1^cKO^ females. RNA concentration and integrity were assessed with Agilent RNA ScreenTape analysis using the Agilent 2100 Bioanalyzer. All samples used in downstream analyses had a RIN >9.7.

#### RNA Sequencing and Analysis

RNA samples were sent to the UCLA Neuroscience Genomics Core for library preparation and sequencing. Sample libraries were generated using TruSeq Stranded RNA kit and sequenced with paired end 150 base pair reads on two lanes using the Illumina HiSeq 3000. Each sample contained between 58-91 million reads (average 79 million reads). All samples raw sequence data passed quality control using FastQC (Andrews, 2010). The data were then mapped to the mouse MM10 genome (Gencode version 17) using STAR aligner (Dobin et al., 2013) with default parameters. MultiQC was used to aggregate quality metrics produced by STAR (Ewels et al., 2016). Within each sample 92-93% of reads were uniquely mapped and used for further analyses. BAM files produced by STAR were sorted and converted to SAM files using samtools (Li et al., 2009). Gene read counts were estimated using HTSeq union gene counts (Anders et al., 2015). Sequencing bias was estimated using Picard Tools (http://picard.sourceforge.net) functions CollectRnaSeqMetrics, CollectGcBiasMetrics, CollectAlignmentSummaryMetrics, and CollectInsertSizeMetrics. An ANOVA was used to determine if these metrics were having a significant effect on the data. Genes with expression (<10 counts) across half the samples were excluded leaving 17,068 genes that passed the filter for further analyses. Principle component analysis was performed in R using the function “prcomp” on data normalized by variance stabilizing transformation (VST). Differential gene expression analysis was performed using DESeq2 (Love et al., 2014) with the model ~Group + Litter. False discovery rate (FDR) < 5% was used as a cutoff to determine if genes were differentially expressed. Gene ontology and pathway analysis were performed using gprofiler2 (https://biit.cs.ut.ee/gprofiler/gost) and disgenet (https://www.disgenet.org/). This work used computational and storage services associated with the Hoffman2 Shared Cluster provided by UCLA Institute for Digital Research and Education’s Research Technology Group. All data and code used in analyses will be shared publicly on GEO and github.

#### Quantitative PCR

Reverse Transcriptase qPCR (RT-qPCR) was performed using the SuperScript IV First-Strand Synthesis System (Invitrogen). For each sample, >500 ng of total RNA was used for cDNA synthesis. In each RT-qPCR reaction, cDNA was combined with LightCycler 480 SYBR Green I

Master Mix (Roche) and exon-spanning primer pairs. All primer pairs were validated for ≥ 1.8 amplification efficiency. Samples were run on a Roche LightCycler 480 real-time PCR system in triplicate, and relative expression levels determined by normalizing the crossing points to the internal reference gene Gapdh. Primers were designed for mouse genes *Slc2a1, Ldha*, and *Vegfa* using Primer3plus (see Key Resources Table for sequences).

#### RNA probe synthesis and *in situ* hybridization

Riboprobes were generated using primers designed against the 5’ or 3’ UTRs of mouse *Aldoa*, *Ldha*, *Slc16a4*, *Pfkl*, *Pfkfb3*, *Pdk1*, *Ccne1*, *Camk2b*, *Dlg4*, *Neo1*, *Sez6*, *Slc2a1*, *Slc2a3*, *Phd2*, *Loxl2*, *Cdh13*, *Igfbp3*, *Vegfr1* and *Vegfa* transcripts. *In situ* hybridization was performed on sections as previously described (Pearson et al., 2011). See Key Resources Table for sequences.

#### RNAScope Fluorescent in situ hybridization

RNAScope FISH was performed on cryosections as per manufacturers guidelines. RNA Scope probe details in Key Resources Table.

#### Immunohistochemistry

Immunohistochemistry was performed on tissue sections as previously described (Pearson *et al*., 2020). Primary antibodies were used at the following dilutions: Foxp1 1:16,000, Glut1 1:100, Laminin 1:1000, Ldha 1:250, Tbr2 1:500, Vegfa 1:250. Donkey secondary antibodies and DAPI were used at 1:1000. 2uM of Isolectin GS-IB4 from *Griffonia simplicifolia*, Alexa Fluor 647 was diluted with secondary antibodies and incubated on slides for 1 hour at room temperature. Sources listed in Key Resources Table.

#### Image Acquisition

Confocal images were acquired using Zeiss LSM 780, LSM 800 and Olympus FV1000 laser scanning confocal microscopes and processed with Zen blue and Fluoview software. DIC imaging of in situ hybridizations were collected using a Zeiss Axioimager microscope and Axiovision software.

### QUANTIFICATION AND STATISTICAL ANALYSIS

Images were processed and compiled using Adobe Photoshop with image adjustments applied to the entire image and restricted to brightness, contrast and levels. Images shown in figures as comparisons, e.g., intensity levels, were obtained and processed in parallel using identical settings. Composite images were assembled using Adobe Illustrator software.

For each experiment, the mean grey values (relative to background) of labeled lateral cortex cells per section was quantified from 3-6 sections per embryo (sampled at ~100μm intervals along the rostro-caudal axis within the presumptive somatosensory cortex). Mean grey value of staining in each mutant was normalized to littermate controls. Mean grey values were measured using Fiji software. For vessel counts and intermediate progenitor counts in Figure 1, contiguous laminin^+^vessels and Tbr2^+^ intermediate progenitors, were counted within a 200μm region in each hemisphere of the lateral cortex (sampled at ~100μm intervals along the rostro-caudal axis within the presumptive somatosensory cortex) for each timepoint. For Figure 4, contiguous vessels in 50μm thick sections were identified using surface rendering software in Imaris. Contiguous vessels were individually color-labelled in Imaris. Each hemisphere was divided into 3 identically sized dorsal-ventral bins (Figure 4E) and vessels within in each bin were quantified. Vessel length and branch points were quantified with Imaris.

gprolifer and disgenet online database tools were used to analyze differentially expressed RNA Sequencing gene lists. Imaris was used to analyze blood vessel length and branching; surface rendered images were generated to identify and measure individual vessels. Graphpad Prism software was used to determine normality of each dataset (using both Shapiro-Wilk test and Kolmogorov-Smirnov Normality tests) and the appropriate parametric test was applied. Student’s t tests and Mann Whitney tests were calculated using Prism software. Significance was assumed when p<0.05. The results of statistical tests (p values and sample sizes) are reported in Figure legends (Student’s t tests were performed unless otherwise stated). Signifiers used are as follows: p > 0.05, ns; * p ≤0.05, ** p ≤0.01, *** p ≤0.001, **** p ≤0.0001. All data are presented as mean ±SEM.

### KEY RESOURCES TABLE

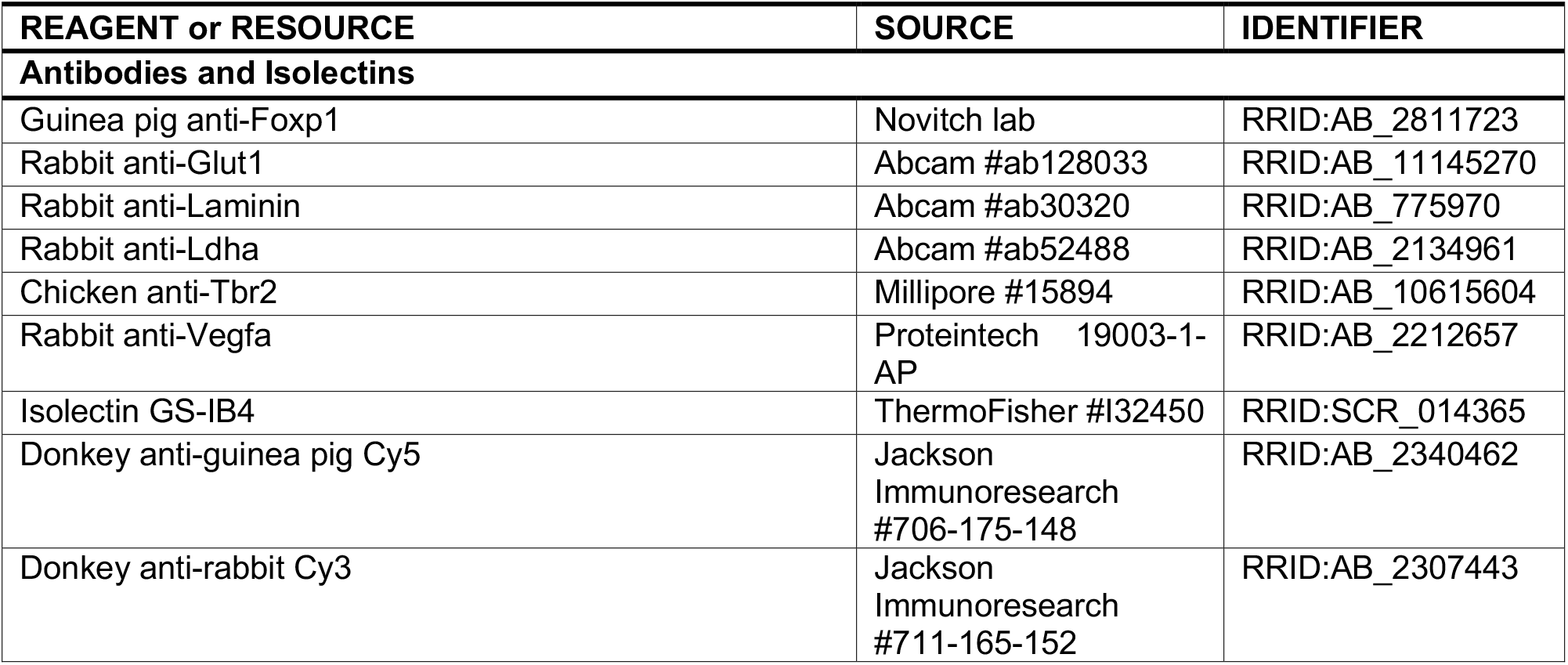
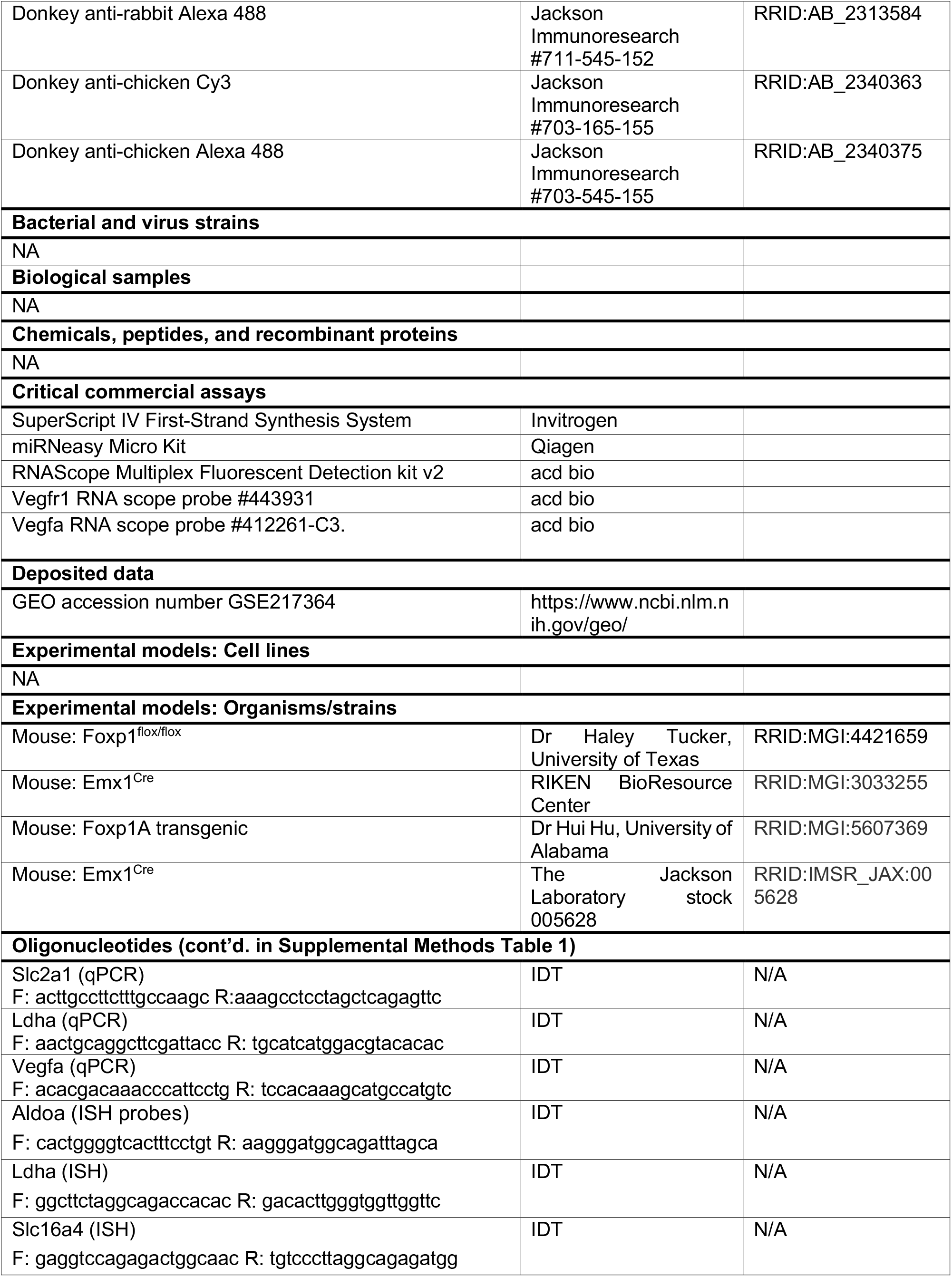
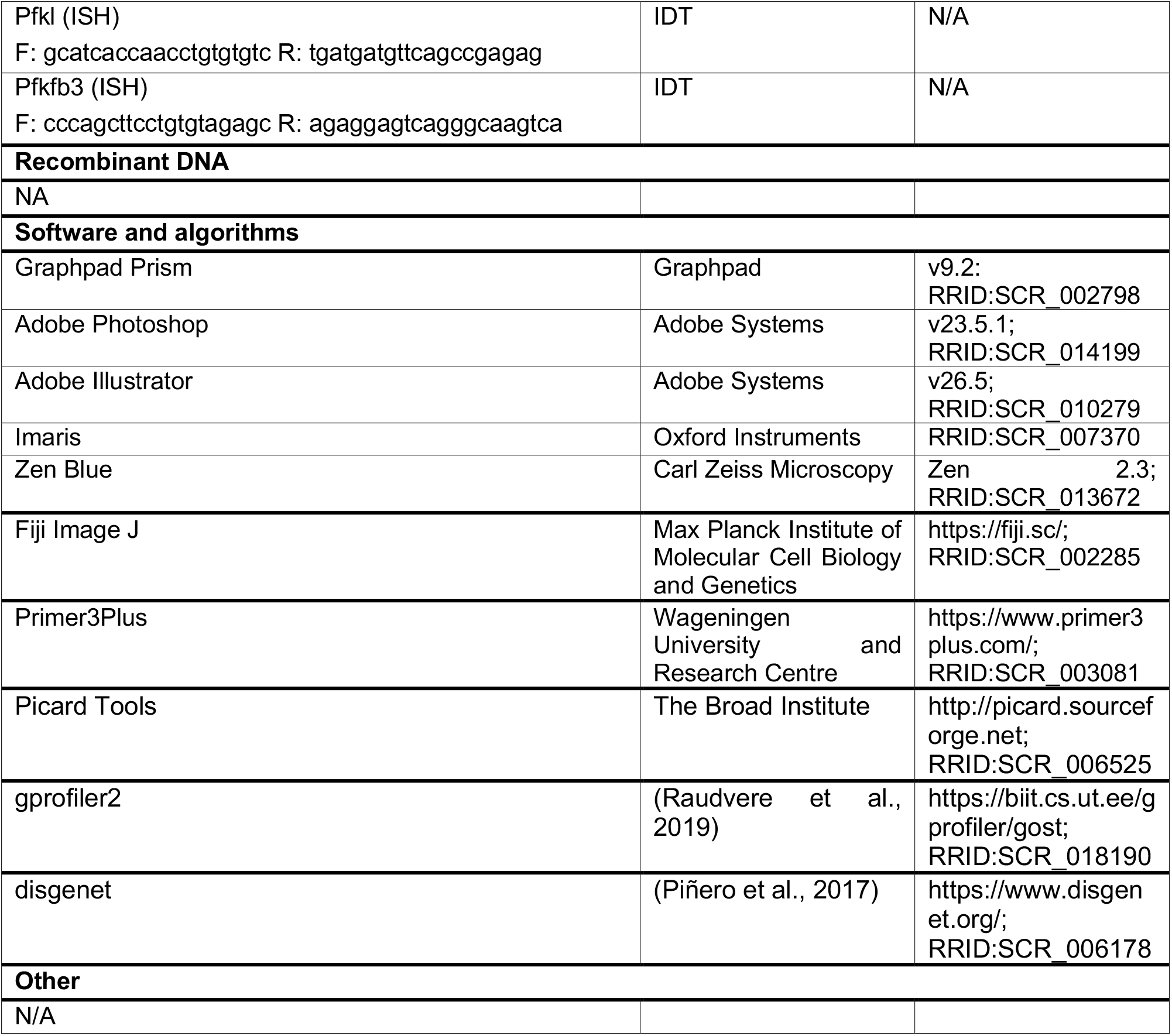

## ACKNOWLEDGEMENTS

We are grateful to H. Tucker, H. Hu, and the RIKEN BioResource Center for mouse strains used in this study. We appreciate K. Phan, A. Schlusche and S. Singh for technical assistance, the UCLA Broad Stem Cell Research Center (BSCRC) for microscopy and other resources, and the UCLA Sequencing Core for technical assistance. We thank G. Manfredi, members of the Novitch lab and the Ross lab for their valuable input and discussions, and M. Placzek, M. Andrews, C. Iadecola, and S. Butler for critical input on the manuscript. This work was supported by research awards from the UCLA Broad Stem Cell Research Center and grants from the NIH R01NS072804 to BGN and R01NS105477 to MER. CAP was supported by a UCLA-California Institute for Regenerative Medicine Training Grant (TG2-01169). J.E.B. was supported by the UCLA BSCRC Rose Hills Foundation Graduate Scholarship Training Program (EDUC4-12753). We also acknowledge the support of the NINDS Informatics Center for Neurogenetics and Neurogenomics (P30NS062691) and the Genetics and Genomics Core of the Semel Institute of Neuroscience at UCLA supported by the NICHD (U54HD087101 and P50HD103557).

## AUTHOR CONTRIBUTIONS

Conceptualization, CAP and BGN. Methodology, CAP. Investigation, CAP, JB, MRMH. Writing, CAP. Review and editing, CAP, MRMH, MER, BGN. Visualization, CAP. Supervision, CAP, MER and BGN. Project administration, CAP. Funding acquisition, CAP, MER, and BGN, Further information and requests for resources and reagents should be directed to the Lead Contact, Caroline Alayne Pearson. This study did not generate unique reagents.

## DECLARATION OF INTERESTS

The authors declare no competing interests.

## Notes

### Competing Interest Statement

The authors have declared no competing interest.

### Summary of Updates

We have revised the submission to adhere to journal formatting guidelines. Supplemental files have been included.

