## Supplemental Figures for "Foxp1 acts upstream of Vegfa, suppresses cortical angiogenesis, and promotes hypoxia in radial glia"

**SUPPLEMENTAL DATA**

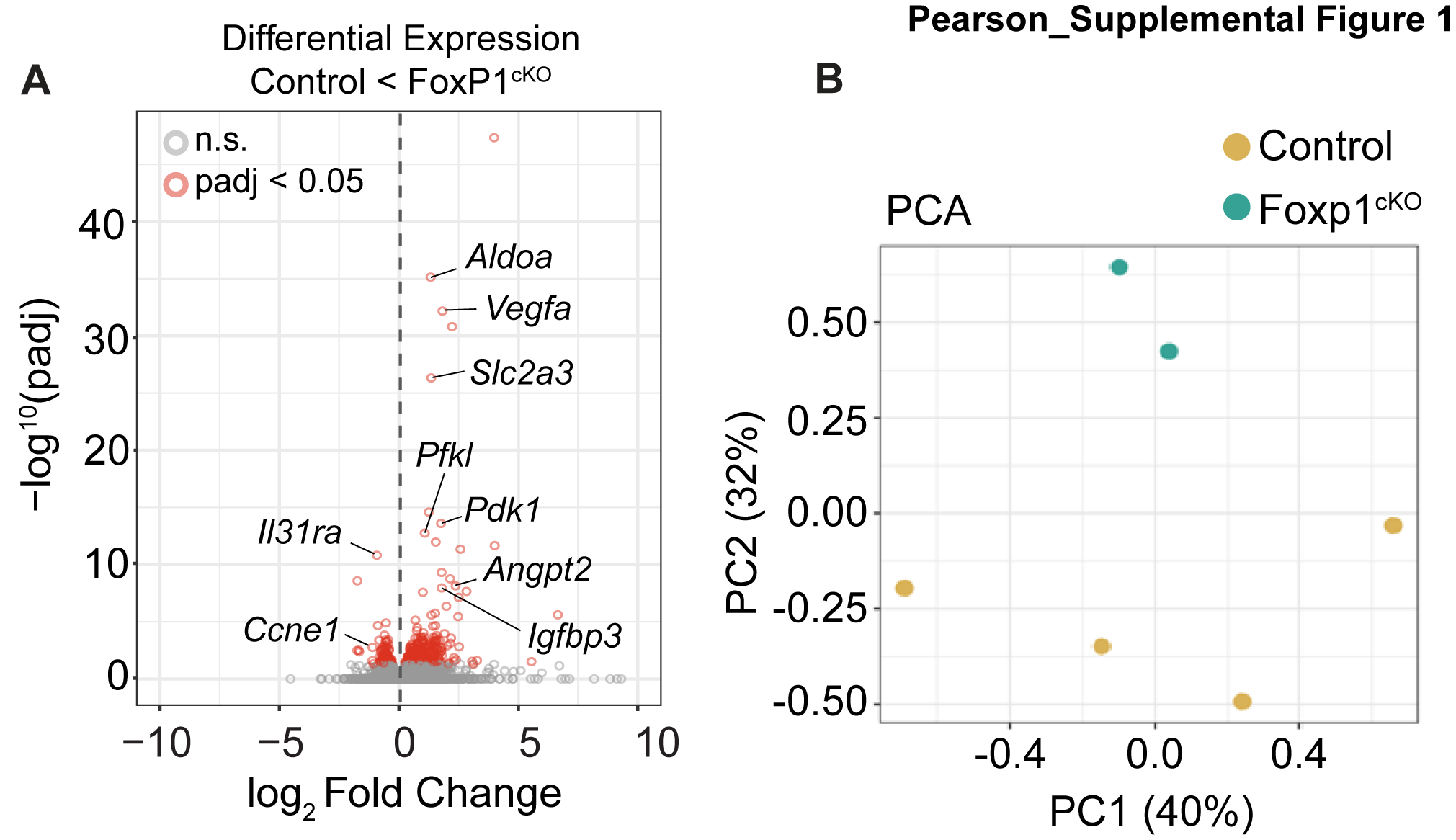

**Figure S1. RNA Seq analysis of Foxp1** **^cKO^ cortex, related to Figure 1.**

(A) Volcano plot of gene expression changes in the absence of Foxp1 in E12.5 lateral cortex compared to WT embryos. Grey circles denote non-significant gene changes (adjusted p-value > 0.05); red circles denote significantly differentially expressed genes (adjusted p-value < 0.05). (B) Principal component analysis (PCA) of control and Foxp1^cKO^ mutants, showing PC2 and PC3.

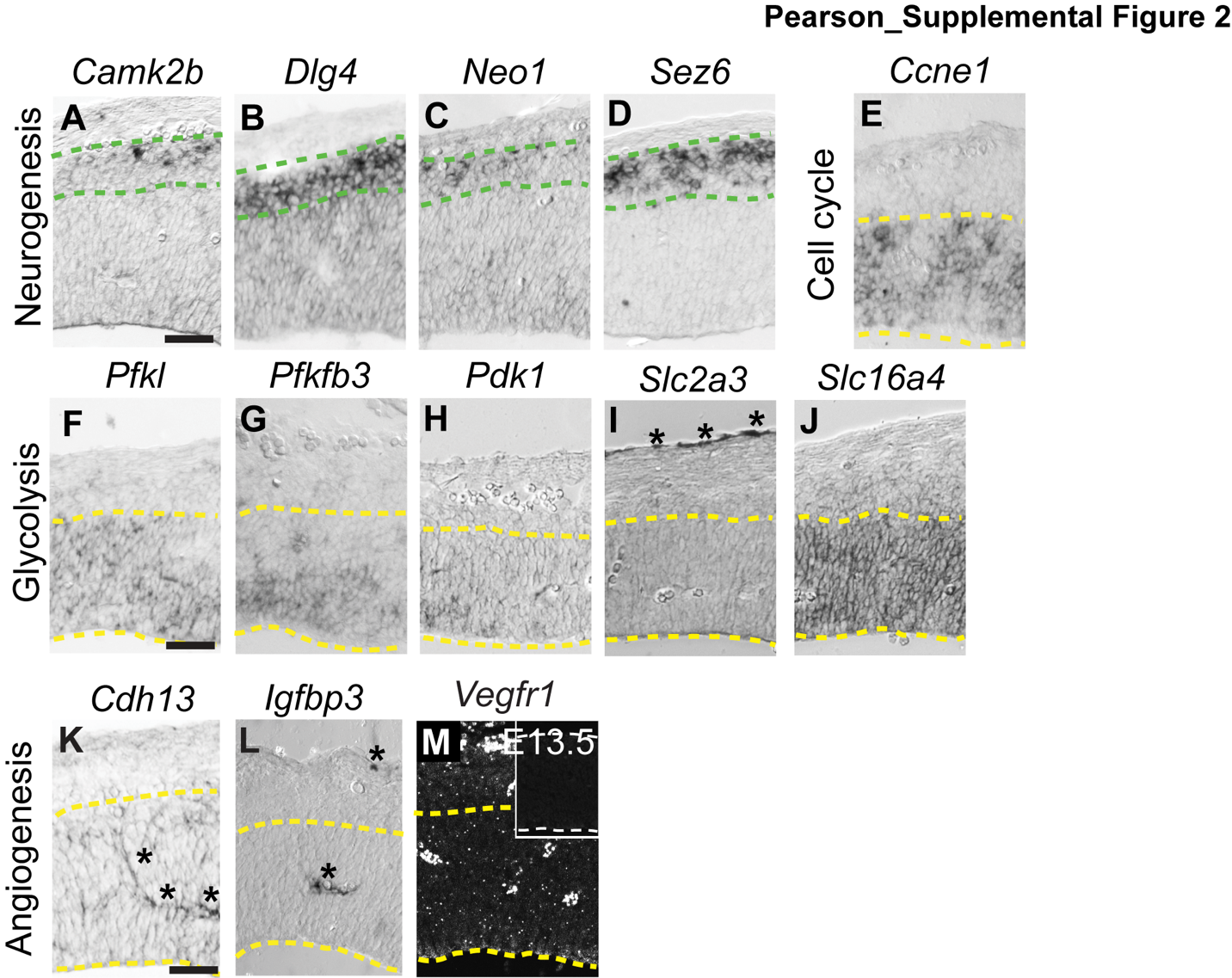

**Figure S2. Wildtype expression of neurogenesis, cell cycle, glycolysis, and angiogenesis genes in the lateral cortex, related to Figure 2.**

(A-D) Wildtype mRNA expression of neurogenesis genes *Camk2b, Dlg4, Neo1* and *Sez6* in the cortical plate and (E) cell cycle gene *Ccne1* in the VZ at E12.5. (F-J) Expression of glycolysis genes Pfkl, Pfkfb3, Pdk1, Slc2a3 and Slc16a4 in the wildtype lateral cortex at E12.5. (K-L) Angiogenesis gene expression including *Cdh13* and *Igfbp3* in the wildtype lateral cortex at E12.5. (M) RNA-Scope FISH analysis of *Vegfr1* mRNA expression in the VZ at E13.5. Negative control for RNA Scope in Opal 690 (inset M). Green dashed lines demarcate the cortical plate; yellow dashed lines demarcate the VZ. Asterisks label endothelial cells. Scale bars 50μm.

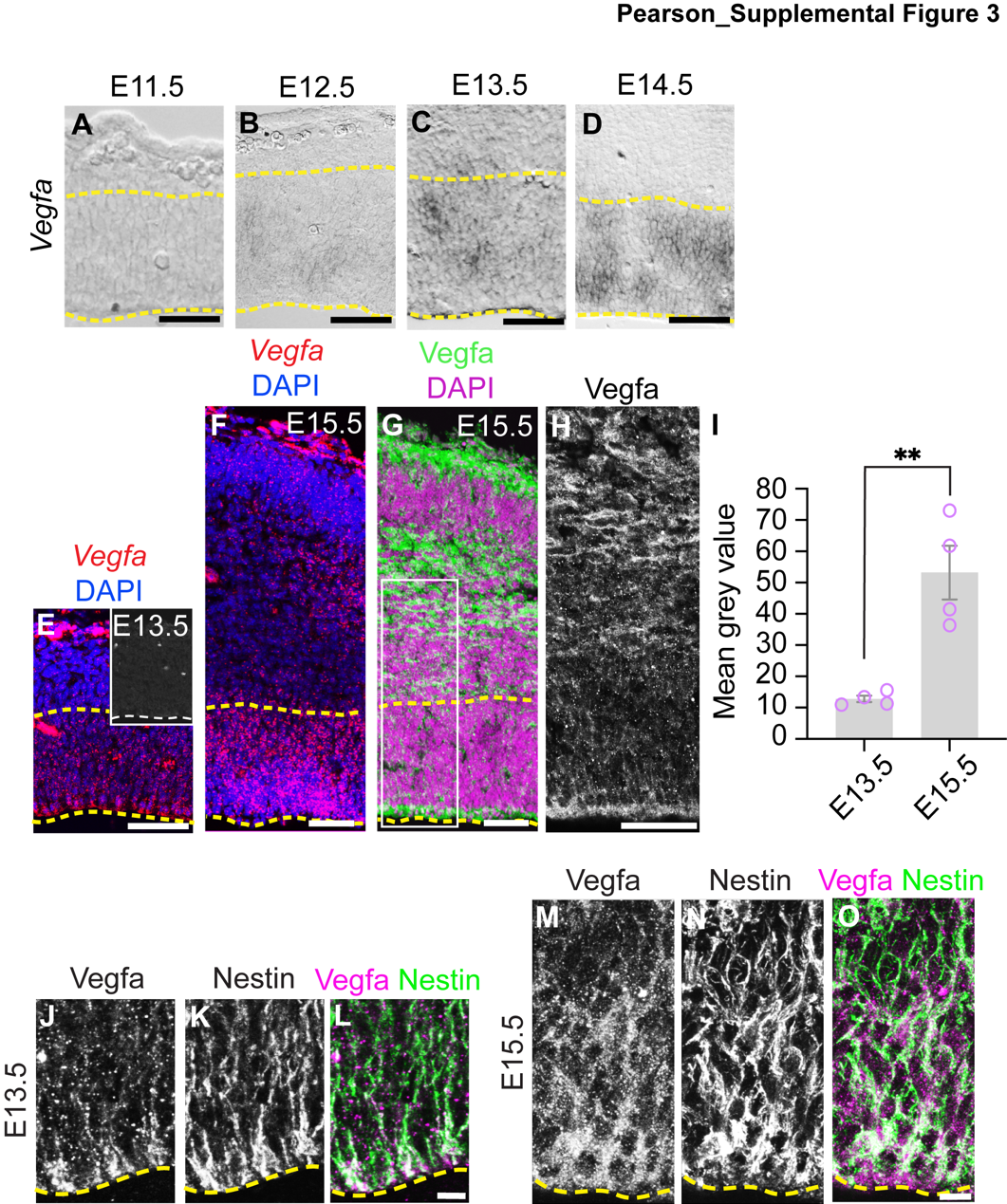

**Figure S3. Vegfa expression increases during neurogenesis, related to Figures 2 and 3.**

(A-D) Wildtype mRNA expression of *Vegfa* in the VZ of the lateral cortex from E11.5 to E14.5. (E-F) RNA Scope FISH analysis of *Vegfa* mRNA expression in the lateral cortex at E13.5 and E15.5. Inset of Opal 520 negative control. (G) IHC for Vegfa in the lateral cortex at E15.5. Inset indicates area magnified in (H). (I) Comparative quantification of *Vegfa* FISH mean grey value in the VZ at E13.5 and E15.5. N= 4 embryos per time point, p**=0.0035. (J-O) High magnification images of the VZ showing Vegfa and Nestin protein expression at E13.5 and E15.5. Scale bars 50μm (A-H) 10μm (J-O).

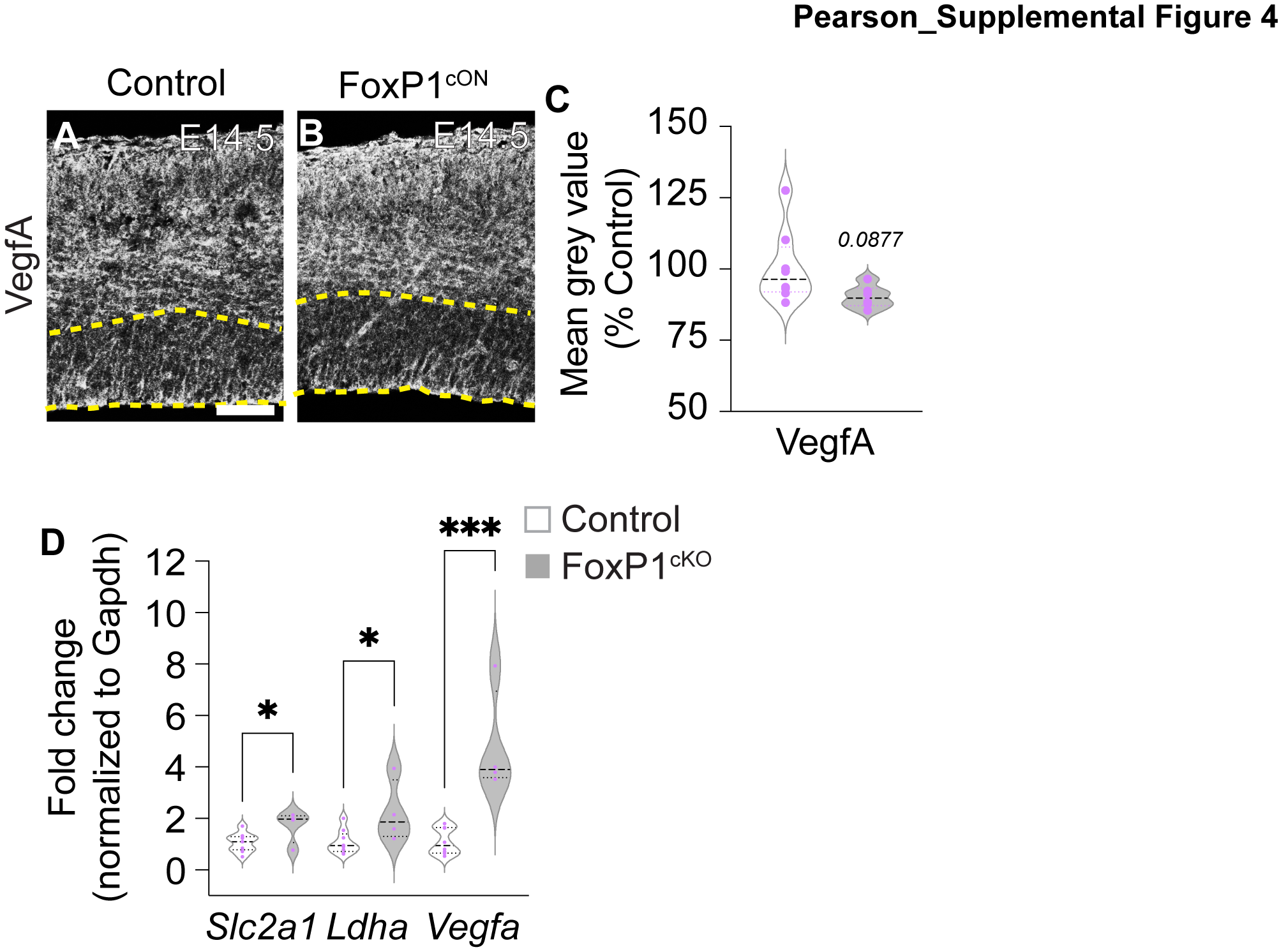

**Figure S4. Vegfa expression in mid-neurogenic Foxp1^ON^ cortex and gene expression changes in the absence of Foxp1, related to Figure 3.**

(A-B) IHC for VegfA in control and Foxp1^cKO^ lateral cortex at E14.5. (C) Quantification of mean gray value of Vegfa expression in control and Foxp1^cKO^ lateral cortex at E14.5. N=2 litters, 8 control, 6 mutants. (D) Quantification of mRNA fold enrichment (normalized to Gapdh) for *Slc2a1, Ldha, Vegfa* in control and Foxp1^cKO^ by qPCR in lateral cortex at E12.5. N = 2 litters, 9 controls, 4 mutants, *p=0.0358, 0.0255 respectively, ***p=0.0006. Scale bar 50μm.

**SUPPLEMENTAL TABLES**

**Table S1. Oligonucleotide sequences for ISH RNA probes (continued from Key Resources table).**

| **Oligonucleotides** | | |
| --- | --- | --- |
| Pdk1 (ISH)  F: ttctcctgcagcctacccta R: gcacccttgtctgagctttc | IDT | N/A |
| Ccne1 (ISH)  F: attgcctccaaagacagtgg R: ctgttggctgacagtggaga | IDT | N/A |
| Camk2b (ISH)  F: ggctttgactgcagaggttc R: caaaagaattgtcccccaga | IDT | N/A |
| Dlg4 (ISH)  F: ctggactgaatagcccaagc R: cccaaaaaccacctttgaga | IDT | N/A |
| Neo1 (ISH)  F: gcaagctctgaaccatcaca R: cctcttccctggacacaaaa | IDT | N/A |
| Sez6 (ISH)  F: gctgtccctcacctcctgta R: cagggcagaaggcttaactg | IDT | N/A |
| Slc2a1 (ISH)  F: agcagtgaagtccaggagga R: ctggtctcaggcaaggaaag | IDT | N/A |
| Slc2a3 (ISH)  F: ccttggctctgctacacaca R: aagtggaggaggggacagtt | IDT | N/A |
| Phd2 (ISH)  F: caagtagagccgtgtgtgga R: ggggttaacggtcctgattt | IDT | N/A |
| Loxl2 (ISH)  F: cgctctggtcttccaaactc R: gcccaccctagacaccagta | IDT | N/A |
| Chd13 (ISH)  F: ctggatttgccctgaatgtt R: catggcatctggtggtactg | IDT | N/A |
| Igfbp3 (ISH)  F: caggaaagaggagtggcttg R: aagctgataggcaggcagaa | IDT | N/A |
| Vegfr1 (ISH)  F: ttctcctgcagcctacccta R: gcacccttgtctgagctttc | IDT | N/A |
| Vegfa (ISH)  F:Gccctgagtcaagaggacag R: ggaagggaagatgaggaagg | IDT | N/A |

**Table S2. Top 80 upregulated genes in Foxp1cKO cortex at E12.5, related to Figures 1 and S1.**

| **Gene Name (Mus musculus)** | **Fold change** | **P value** | **Description** |
| --- | --- | --- | --- |
| Ddn | 6.65539649 | 4.34E-09 | dendrin [Source:MGI Symbol;Acc:MGI:108101] |
| Gm29683 | 5.55131856 | 0.00081083 | predicted gene, 29683 [Source:MGI Symbol;Acc:MGI:5588842] |
| Gm15283 | 4.01183319 | 1.66E-15 | predicted gene 15283 [Source:MGI Symbol;Acc:MGI:3705161] |
| F630040K05Rik | 3.98810861 | 3.55E-52 | RIKEN cDNA F630040K05 gene [Source:MGI Symbol;Acc:MGI:4437734] |
| Rasgef1a | 3.27179303 | 0.00053465 | RasGEF domain family, member 1A [Source:MGI Symbol;Acc:MGI:1917977] |
| CT010460.2 | 3.11608167 | 0.00193561 |  |
| Fam19a1 | 3.0666215 | 0.00070148 | family with sequence similarity 19, member A1 [Source:MGI Symbol;Acc:MGI:2443695] |
| Esm1 | 2.829631 | 3.05E-11 | endothelial cell-specific molecule 1 [Source:MGI Symbol;Acc:MGI:1918940] |
| Kcne3 | 2.57756205 | 3.81E-15 | potassium voltage-gated channel, Isk-related subfamily, gene 3 [Source:MGI Symbol;Acc:MGI:1891124] |
| Mctp1 | 2.51954838 | 9.53E-06 | multiple C2 domains, transmembrane 1 [Source:MGI Symbol;Acc:MGI:1926021] |
| Adm | 2.50655921 | 1.12E-10 | adrenomedullin [Source:MGI Symbol;Acc:MGI:108058] |
| Ciart | 2.48310475 | 6.80E-09 | circadian associated repressor of transcription [Source:MGI Symbol;Acc:MGI:2684975] |
| Egr3 | 2.39720897 | 0.00041827 | early growth response 3 [Source:MGI Symbol;Acc:MGI:1306780] |
| Angpt2 | 2.37654774 | 8.90E-12 | angiopoietin 2 [Source:MGI Symbol;Acc:MGI:1202890] |
| Cntnap1 | 2.30991797 | 0.00022566 | contactin associated protein-like 1 [Source:MGI Symbol;Acc:MGI:1858201] |
| Caln1 | 2.29886844 | 0.00070511 | calneuron 1 [Source:MGI Symbol;Acc:MGI:2155987] |
| Bnip3 | 2.22446793 | 4.69E-35 | BCL2/adenovirus E1B interacting protein 3 [Source:MGI Symbol;Acc:MGI:109326] |
| Rps3a2 | 2.18806411 | 8.52E-07 | ribosomal protein S3A2 [Source:MGI Symbol;Acc:MGI:3642853] |
| Serpine1 | 2.15823855 | 3.73E-06 | serine (or cysteine) peptidase inhibitor, clade E, member 1 [Source:MGI Symbol;Acc:MGI:97608] |
| Slc16a3 | 2.1459402 | 1.88E-12 | solute carrier family 16 (monocarboxylic acid transporters), member 3 [Source:MGI Symbol;Acc:MGI:1933438] |
| Hpcal4 | 2.07681771 | 0.00122888 | hippocalcin-like 4 [Source:MGI Symbol;Acc:MGI:2157521] |
| Otof | 2.07208059 | 0.00195399 | otoferlin [Source:MGI Symbol;Acc:MGI:1891247] |
| Plppr4 | 2.02429672 | 7.38E-06 | phospholipid phosphatase related 4 [Source:MGI Symbol;Acc:MGI:106530] |
| 8430408G22Rik | 1.98774221 | 7.16E-10 | RIKEN cDNA 8430408G22 gene [Source:MGI Symbol;Acc:MGI:1918730] |
| Ackr2 | 1.90982731 | 2.87E-07 | atypical chemokine receptor 2 [Source:MGI Symbol;Acc:MGI:1891697] |
| Vegfa | 1.82627743 | 1.54E-36 | vascular endothelial growth factor A [Source:MGI Symbol;Acc:MGI:103178] |
| P4ha2 | 1.79677903 | 0.00012164 | procollagen-proline, 2-oxoglutarate 4-dioxygenase (proline 4-hydroxylase), alpha II polypeptide [Source:MGI Symbol;Acc:MGI:894286] |
| Igfbp3 | 1.79406457 | 1.44E-11 | insulin-like growth factor binding protein 3 [Source:MGI Symbol;Acc:MGI:96438] |
| Grm5 | 1.79178696 | 1.11E-05 | glutamate receptor, metabotropic 5 [Source:MGI Symbol;Acc:MGI:1351342] |
| Pfkp | 1.78998445 | 4.81E-13 | phosphofructokinase, platelet [Source:MGI Symbol;Acc:MGI:1891833] |
| Rgs4 | 1.78474484 | 3.20E-05 | regulator of G-protein signaling 4 [Source:MGI Symbol;Acc:MGI:108409] |
| Pdk1 | 1.76402988 | 1.35E-17 | pyruvate dehydrogenase kinase, isoenzyme 1 [Source:MGI Symbol;Acc:MGI:1926119] |
| Stc2 | 1.76397292 | 4.01E-05 | stanniocalcin 2 [Source:MGI Symbol;Acc:MGI:1316731] |
| Ier3 | 1.76376615 | 1.49E-05 | immediate early response 3 [Source:MGI Symbol;Acc:MGI:104814] |
| Coch | 1.73031956 | 0.00067348 | cochlin [Source:MGI Symbol;Acc:MGI:1278313] |
| A2m | 1.70106531 | 1.59E-06 | alpha-2-macroglobulin [Source:MGI Symbol;Acc:MGI:2449119] |
| Cacna1i | 1.68718652 | 0.00097619 | calcium channel, voltage-dependent, alpha 1I subunit [Source:MGI Symbol;Acc:MGI:2178051] |
| Sptbn4 | 1.68218143 | 7.86E-05 | spectrin beta, non-erythrocytic 4 [Source:MGI Symbol;Acc:MGI:1890574] |
| Gpr158 | 1.66897042 | 0.00028633 | G protein-coupled receptor 158 [Source:MGI Symbol;Acc:MGI:2441697] |
| Spock3 | 1.6566705 | 0.00053719 | sparc/osteonectin, cwcv and kazal-like domains proteoglycan 3 [Source:MGI Symbol;Acc:MGI:1920152] |
| Gm996 | 1.65486028 | 3.27E-05 | predicted gene 996 [Source:MGI Symbol;Acc:MGI:2685842] |
| Caly | 1.65031118 | 0.00020828 | calcyon neuron-specific vesicular protein [Source:MGI Symbol;Acc:MGI:1915816] |
| Faim2 | 1.63361551 | 1.74E-05 | Fas apoptotic inhibitory molecule 2 [Source:MGI Symbol;Acc:MGI:1919643] |
| Plppr5 | 1.61519519 | 1.47E-06 | phospholipid phosphatase related 5 [Source:MGI Symbol;Acc:MGI:1923019] |
| Lgi1 | 1.60916945 | 2.20E-05 | leucine-rich repeat LGI family, member 1 [Source:MGI Symbol;Acc:MGI:1861691] |
| Ptprn | 1.60844122 | 0.00102085 | protein tyrosine phosphatase, receptor type, N [Source:MGI Symbol;Acc:MGI:102765] |
| Grin1 | 1.57363908 | 1.11E-05 | glutamate receptor, ionotropic, NMDA1 (zeta 1) [Source:MGI Symbol;Acc:MGI:95819] |
| Tfap2d | 1.57185471 | 4.32E-07 | transcription factor AP-2, delta [Source:MGI Symbol;Acc:MGI:2153466] |
| Gipr | 1.55868747 | 1.49E-06 | gastric inhibitory polypeptide receptor [Source:MGI Symbol;Acc:MGI:1352753] |
| Gt(ROSA)26Sor | 1.54378507 | 7.36E-16 | gene trap ROSA 26, Philippe Soriano [Source:MGI Symbol;Acc:MGI:104735] |
| Ppp1r14c | 1.53532665 | 2.29E-06 | protein phosphatase 1, regulatory (inhibitor) subunit 14c [Source:MGI Symbol;Acc:MGI:1923392] |
| Pitpnm3 | 1.53304006 | 0.00089264 | PITPNM family member 3 [Source:MGI Symbol;Acc:MGI:2685726] |
| Vldlr | 1.5304937 | 4.82E-08 | very low density lipoprotein receptor [Source:MGI Symbol;Acc:MGI:98935] |
| Ddit4 | 1.52883808 | 7.82E-07 | DNA-damage-inducible transcript 4 [Source:MGI Symbol;Acc:MGI:1921997] |
| Grin2b | 1.52478075 | 5.74E-05 | glutamate receptor, ionotropic, NMDA2B (epsilon 2) [Source:MGI Symbol;Acc:MGI:95821] |
| Fstl4 | 1.52292707 | 0.00011112 | follistatin-like 4 [Source:MGI Symbol;Acc:MGI:2443199] |
| Tnfaip3 | 1.52240613 | 9.99E-06 | tumor necrosis factor, alpha-induced protein 3 [Source:MGI Symbol;Acc:MGI:1196377] |
| Flt1 | 1.52183643 | 3.06E-09 | FMS-like tyrosine kinase 1 [Source:MGI Symbol;Acc:MGI:95558] |
| Arpp21 | 1.51578087 | 2.12E-06 | cyclic AMP-regulated phosphoprotein, 21 [Source:MGI Symbol;Acc:MGI:107562] |
| Pfkfb3 | 1.50878807 | 3.17E-06 | 6-phosphofructo-2-kinase/fructose-2,6-biphosphatase 3 [Source:MGI Symbol;Acc:MGI:2181202] |
| Ndufa4l2 | 1.48736539 | 0.00014423 | NADH dehydrogenase (ubiquinone) 1 alpha subcomplex, 4-like 2 [Source:MGI Symbol;Acc:MGI:3039567] |
| Ablim3 | 1.47532395 | 0.00069143 | actin binding LIM protein family, member 3 [Source:MGI Symbol;Acc:MGI:2442582] |
| C1ql3 | 1.45828294 | 0.00160106 | C1q-like 3 [Source:MGI Symbol;Acc:MGI:2387350] |
| Unc80 | 1.45537429 | 0.00113367 | unc-80, NALCN activator [Source:MGI Symbol;Acc:MGI:2652882] |
| Pde8b | 1.44773377 | 4.92E-06 | phosphodiesterase 8B [Source:MGI Symbol;Acc:MGI:2443999] |
| Aldh1l1 | 1.4451944 | 0.00036134 | aldehyde dehydrogenase 1 family, member L1 [Source:MGI Symbol;Acc:MGI:1340024] |
| Camk2b | 1.43892756 | 7.04E-07 | calcium/calmodulin-dependent protein kinase II, beta [Source:MGI Symbol;Acc:MGI:88257] |
| Dync1i1 | 1.43545093 | 1.85E-06 | dynein cytoplasmic 1 intermediate chain 1 [Source:MGI Symbol;Acc:MGI:107743] |
| Lhfpl3 | 1.43425078 | 0.0008372 | lipoma HMGIC fusion partner-like 3 [Source:MGI Symbol;Acc:MGI:1925076] |
| Rimbp2 | 1.43058043 | 0.00137117 | RIMS binding protein 2 [Source:MGI Symbol;Acc:MGI:2443235] |
| Prelid2 | 1.4273582 | 5.63E-08 | PRELI domain containing 2 [Source:MGI Symbol;Acc:MGI:1924869] |
| Scg2 | 1.42037754 | 1.98E-05 | secretogranin II [Source:MGI Symbol;Acc:MGI:103033] |
| Ntsr1 | 1.41471679 | 0.00025169 | neurotensin receptor 1 [Source:MGI Symbol;Acc:MGI:97386] |
| Eef1a2 | 1.40776743 | 6.59E-06 | eukaryotic translation elongation factor 1 alpha 2 [Source:MGI Symbol;Acc:MGI:1096317] |
| Egln3 | 1.39567427 | 4.49E-05 | egl-9 family hypoxia-inducible factor 3 [Source:MGI Symbol;Acc:MGI:1932288] |
| Ndrg1 | 1.3872471 | 0.00018288 | N-myc downstream regulated gene 1 [Source:MGI Symbol;Acc:MGI:1341799] |
| Adap1 | 1.38600031 | 0.00100473 | ArfGAP with dual PH domains 1 [Source:MGI Symbol;Acc:MGI:2442201] |
| Rgs6 | 1.38567624 | 0.00068082 | regulator of G-protein signaling 6 [Source:MGI Symbol;Acc:MGI:1354730] |
| P4ha1 | 1.36681882 | 4.75E-09 | procollagen-proline, 2-oxoglutarate 4-dioxygenase (proline 4-hydroxylase), alpha 1 polypeptide [Source:MGI Symbol;Acc:MGI:97463] |
| Adora2a | 1.36165565 | 0.00130986 | adenosine A2a receptor [Source:MGI Symbol;Acc:MGI:99402] |

**Table S3. Top 80 downregulated genes in Foxp1cKO cortex at E12.5, related to Figures 1 and S1.**

| ***Gene Name (Mus musculus)*** | ***Fold change*** | ***P value*** | ***Description*** |
| --- | --- | --- | --- |
| Gm21887 | -1.7454143 | 2.93E-05 | predicted gene, 21887 [Source:MGI Symbol;Acc:MGI:5434051] |
| Txnip | -1.73318 | 3.03E-12 | thioredoxin interacting protein [Source:MGI Symbol;Acc:MGI:1889549] |
| C030037D09Rik | -1.6809915 | 2.74E-05 | RIKEN cDNA C030037D09 gene [Source:MGI Symbol;Acc:MGI:1924865] |
| AA465934 | -1.6576161 | 4.33E-05 | expressed sequence AA465934 [Source:MGI Symbol;Acc:MGI:2671018] |
| 9930014A18Rik | -1.1108228 | 0.00056841 | RIKEN cDNA 9930014A18 gene [Source:MGI Symbol;Acc:MGI:2444091] |
| Ccne1 | -1.1039293 | 1.22E-05 | cyclin E1 [Source:MGI Symbol;Acc:MGI:88316] |
| 1600010M07Rik | -1.0650143 | 0.00184514 | RIKEN cDNA 1600010M07 gene [Source:MGI Symbol;Acc:MGI:1917031] |
| Il31ra | -0.9216888 | 1.40E-14 | interleukin 31 receptor A [Source:MGI Symbol;Acc:MGI:2180511] |
| Snapc5 | -0.8809404 | 4.82E-08 | small nuclear RNA activating complex, polypeptide 5 [Source:MGI Symbol;Acc:MGI:1914282] |
| Gm15564 | -0.8262105 | 1.63E-06 | predicted gene 15564 [Source:MGI Symbol;Acc:MGI:3783013] |
| Rpp25l | -0.8001076 | 0.0001774 | ribonuclease P/MRP 25 subunit-like [Source:MGI Symbol;Acc:MGI:1917211] |
| Gm4430 | -0.7812544 | 0.00072593 | predicted gene 4430 [Source:MGI Symbol;Acc:MGI:3782614] |
| 4930427A07Rik | -0.7579571 | 4.38E-05 | RIKEN cDNA 4930427A07 gene [Source:MGI Symbol;Acc:MGI:2144738] |
| Dhrs4 | -0.7578561 | 0.00076148 | dehydrogenase/reductase (SDR family) member 4 [Source:MGI Symbol;Acc:MGI:90169] |
| Rfc4 | -0.7527353 | 2.59E-05 | replication factor C (activator 1) 4 [Source:MGI Symbol;Acc:MGI:2146571] |
| Mcur1 | -0.7456268 | 0.00075977 | mitochondrial calcium uniporter regulator 1 [Source:MGI Symbol;Acc:MGI:1923387] |
| Ung | -0.7436174 | 0.00051431 | uracil DNA glycosylase [Source:MGI Symbol;Acc:MGI:109352] |
| Rps3a1 | -0.7167743 | 0.00015373 | ribosomal protein S3A1 [Source:MGI Symbol;Acc:MGI:1202063] |
| Slc25a10 | -0.7167095 | 2.32E-05 | solute carrier family 25 (mitochondrial carrier, dicarboxylate transporter), member 10 [Source:MGI Symbol;Acc:MGI:1353497] |
| Ide | -0.7047055 | 0.00040817 | insulin degrading enzyme [Source:MGI Symbol;Acc:MGI:96412] |
| Rcc1 | -0.6926344 | 0.00028344 | regulator of chromosome condensation 1 [Source:MGI Symbol;Acc:MGI:1913989] |
| 2310011J03Rik | -0.690648 | 0.00195057 | RIKEN cDNA 2310011J03 gene [Source:MGI Symbol;Acc:MGI:1913624] |
| Prdx1 | -0.6842499 | 0.00053835 | peroxiredoxin 1 [Source:MGI Symbol;Acc:MGI:99523] |
| Myb | -0.6799152 | 0.00090893 | myeloblastosis oncogene [Source:MGI Symbol;Acc:MGI:97249] |
| Pno1 | -0.6761335 | 1.35E-05 | partner of NOB1 homolog [Source:MGI Symbol;Acc:MGI:1913499] |
| Tyms | -0.6738263 | 8.02E-05 | thymidylate synthase [Source:MGI Symbol;Acc:MGI:98878] |
| Arrdc4 | -0.6635273 | 2.16E-05 | arrestin domain containing 4 [Source:MGI Symbol;Acc:MGI:1913662] |
| Lsm5 | -0.6616134 | 0.00048587 | LSM5 homolog, U6 small nuclear RNA and mRNA degradation associated [Source:MGI Symbol;Acc:MGI:1913623] |
| Fam72a | -0.6612573 | 0.00018603 | family with sequence similarity 72, member A [Source:MGI Symbol;Acc:MGI:1919669] |
| Amd1 | -0.6491136 | 0.00060853 | S-adenosylmethionine decarboxylase 1 [Source:MGI Symbol;Acc:MGI:88004] |
| Prx | -0.638529 | 0.00076685 | periaxin [Source:MGI Symbol;Acc:MGI:108176] |
| Gcsh | -0.6369482 | 0.00137435 | glycine cleavage system protein H (aminomethyl carrier) [Source:MGI Symbol;Acc:MGI:1915383] |
| Dut | -0.6356397 | 0.00034822 | deoxyuridine triphosphatase [Source:MGI Symbol;Acc:MGI:1346051] |
| AC165092.1 | -0.6350126 | 0.00127199 |  |
| Fanca | -0.6278163 | 0.0012161 | Fanconi anemia, complementation group A [Source:MGI Symbol;Acc:MGI:1341823] |
| Pcna | -0.6278124 | 0.00050695 | proliferating cell nuclear antigen [Source:MGI Symbol;Acc:MGI:97503] |
| Akr1b3 | -0.6113962 | 0.00011939 | aldo-keto reductase family 1, member B3 (aldose reductase) [Source:MGI Symbol;Acc:MGI:1353494] |
| Aurka | -0.6080558 | 0.00152722 | aurora kinase A [Source:MGI Symbol;Acc:MGI:894678] |
| Prokr1 | -0.6043059 | 0.00037827 | prokineticin receptor 1 [Source:MGI Symbol;Acc:MGI:1929676] |
| Ccdc25 | -0.5995813 | 3.90E-07 | coiled-coil domain containing 25 [Source:MGI Symbol;Acc:MGI:1914429] |
| Pclaf | -0.5913014 | 2.02E-05 | PCNA clamp associated factor [Source:MGI Symbol;Acc:MGI:1915276] |
| Timm22 | -0.5798368 | 7.52E-05 | translocase of inner mitochondrial membrane 22 [Source:MGI Symbol;Acc:MGI:1929742] |
| Cycs | -0.578209 | 1.86E-05 | cytochrome c, somatic [Source:MGI Symbol;Acc:MGI:88578] |
| Dhfr | -0.5775575 | 0.00042771 | dihydrofolate reductase [Source:MGI Symbol;Acc:MGI:94890] |
| Utp6 | -0.5743641 | 0.00065969 | UTP6 small subunit processome component [Source:MGI Symbol;Acc:MGI:2445193] |
| Mrpl12 | -0.5672068 | 7.76E-05 | mitochondrial ribosomal protein L12 [Source:MGI Symbol;Acc:MGI:1926273] |
| Tbl3 | -0.5631961 | 0.00135835 | transducin (beta)-like 3 [Source:MGI Symbol;Acc:MGI:2384863] |
| Umps | -0.5599798 | 0.00191753 | uridine monophosphate synthetase [Source:MGI Symbol;Acc:MGI:1298388] |
| Larp7 | -0.5587161 | 7.48E-06 | La ribonucleoprotein domain family, member 7 [Source:MGI Symbol;Acc:MGI:107634] |
| Pole3 | -0.5569267 | 0.00156599 | polymerase (DNA directed), epsilon 3 (p17 subunit) [Source:MGI Symbol;Acc:MGI:1933378] |
| Ahcy | -0.5559686 | 2.52E-06 | S-adenosylhomocysteine hydrolase [Source:MGI Symbol;Acc:MGI:87968] |
| Rrm2 | -0.5538147 | 2.64E-05 | ribonucleotide reductase M2 [Source:MGI Symbol;Acc:MGI:98181] |
| Wdhd1 | -0.5485145 | 0.00115459 | WD repeat and HMG-box DNA binding protein 1 [Source:MGI Symbol;Acc:MGI:2443514] |
| Slbp | -0.5479877 | 0.00032909 | stem-loop binding protein [Source:MGI Symbol;Acc:MGI:108402] |
| Rfc3 | -0.5449692 | 1.38E-06 | replication factor C (activator 1) 3 [Source:MGI Symbol;Acc:MGI:1916513] |
| Cdc6 | -0.5439973 | 0.00125598 | cell division cycle 6 [Source:MGI Symbol;Acc:MGI:1345150] |
| Tubd1 | -0.5430312 | 0.00131465 | tubulin, delta 1 [Source:MGI Symbol;Acc:MGI:1891826] |
| Ppa1 | -0.5374194 | 2.60E-08 | pyrophosphatase (inorganic) 1 [Source:MGI Symbol;Acc:MGI:97831] |
| Spdl1 | -0.5369133 | 5.73E-05 | spindle apparatus coiled-coil protein 1 [Source:MGI Symbol;Acc:MGI:1917635] |
| Limd2 | -0.5353452 | 0.00029489 | LIM domain containing 2 [Source:MGI Symbol;Acc:MGI:1915053] |
| Rrm1 | -0.5334689 | 0.00046298 | ribonucleotide reductase M1 [Source:MGI Symbol;Acc:MGI:98180] |
| 2310033P09Rik | -0.5333676 | 0.00183879 | RIKEN cDNA 2310033P09 gene [Source:MGI Symbol;Acc:MGI:1915112] |
| Atp5k | -0.5296294 | 8.18E-05 | ATP synthase, H+ transporting, mitochondrial F1F0 complex, subunit E [Source:MGI Symbol;Acc:MGI:106636] |
| Capza1 | -0.5262706 | 0.00175806 | capping protein (actin filament) muscle Z-line, alpha 1 [Source:MGI Symbol;Acc:MGI:106227] |
| Prim1 | -0.5248936 | 0.00178259 | DNA primase, p49 subunit [Source:MGI Symbol;Acc:MGI:97757] |
| Itpa | -0.5246876 | 0.00013457 | inosine triphosphatase (nucleoside triphosphate pyrophosphatase) [Source:MGI Symbol;Acc:MGI:96622] |
| Psmc5 | -0.5232555 | 0.00036823 | protease (prosome, macropain) 26S subunit, ATPase 5 [Source:MGI Symbol;Acc:MGI:105047] |
| Hmgn2 | -0.5217042 | 0.00024074 | high mobility group nucleosomal binding domain 2 [Source:MGI Symbol;Acc:MGI:96136] |
| Phf5a | -0.5197565 | 0.00162192 | PHD finger protein 5A [Source:MGI Symbol;Acc:MGI:2156864] |
| Nudcd2 | -0.5191933 | 0.00146025 | NudC domain containing 2 [Source:MGI Symbol;Acc:MGI:1277103] |
| Alyref | -0.5186022 | 1.03E-05 | Aly/REF export factor [Source:MGI Symbol;Acc:MGI:1341044] |
| Tsfm | -0.5173147 | 0.00064066 | Ts translation elongation factor, mitochondrial [Source:MGI Symbol;Acc:MGI:1913649] |
| Heatr1 | -0.5163531 | 0.00069873 | HEAT repeat containing 1 [Source:MGI Symbol;Acc:MGI:2442524] |
| Cenpu | -0.5144973 | 0.00123 | centromere protein U [Source:MGI Symbol;Acc:MGI:1919126] |
| Dnajc9 | -0.5127613 | 0.00166414 | DnaJ heat shock protein family (Hsp40) member C9 [Source:MGI Symbol;Acc:MGI:1915326] |
| Arpc5l | -0.5114661 | 6.27E-05 | actin related protein 2/3 complex, subunit 5-like [Source:MGI Symbol;Acc:MGI:1921442] |
| Pola1 | -0.5094492 | 2.19E-06 | polymerase (DNA directed), alpha 1 [Source:MGI Symbol;Acc:MGI:99660] |
| Ppil1 | -0.5067604 | 9.25E-05 | peptidylprolyl isomerase (cyclophilin)-like 1 [Source:MGI Symbol;Acc:MGI:1916066] |
| Abcf2 | -0.5054141 | 0.00140308 | ATP-binding cassette, sub-family F (GCN20), member 2 [Source:MGI Symbol;Acc:MGI:1351657] |
| Erh | -0.5036326 | 0.00135281 | enhancer of rudimentary homolog (Drosophila) [Source:MGI Symbol;Acc:MGI:108089] |
